# Simultaneous Confidence Regions for Image Excursion Sets: a Validation Study with Applications in fMRI

**DOI:** 10.1101/2025.01.24.634784

**Authors:** Jiyue Qin, Samuel Davenport, Armin Schwartzman

## Abstract

Functional Magnetic Resonance Imaging (fMRI) is commonly used to localize brain regions activated during a task. Methods have been developed for constructing confidence regions of image excursion sets, allowing inference on brain regions exceeding non-zero activation thresholds. However, these methods have been limited to a single predefined threshold and brain volume data, overlooking more sensitive cortical surface analyses. We present an approach that constructs simultaneous confidence regions (SCRs) which are valid for all possible activation thresholds and are applicable to both volume and surface data. This approach is based on a recent method that constructs SCRs from simultaneous confidence bands (SCBs), obtained by using the bootstrap on 1D and 2D images. To extend this method to fMRI studies, we evaluate the validity of the bootstrap with fMRI data through extensive 2D simulations. Six bootstrap variants, including the nonparametric bootstrap and multiplier bootstrap are compared. The Rademacher multiplier bootstrap-t performs the best, achieving a coverage rate close to the nominal level with sample sizes as low as 20. We further validate our approach using realistic noise simulations obtained by resampling resting-state 3D fMRI data, a technique that has become the gold standard in the field. Moreover, our implementation handles data of any dimension and is equipped with interactive visualization tools designed for fMRI analysis. We apply our approach to task fMRI volume data and surface data from the Human Connectome Project, showcasing the method’s utility.

## 1 Introduction

Functional Magnetic Resonance Imaging (fMRI) is a widely used noninvasive neuroimaging technique for measuring brain activity by detecting changes in blood flow (Lindquist, 2008). During an fMRI experiment, a participant undergoes a series of scans while performing a task. Each scan generates a 3D image of the brain, consisting of over 200,000 voxels, where the image intensity at each voxel represents the brain activity at that location (Cremers et al., 2017). A first-level analysis is performed to create a 3D contrast image, which represents the change in brain activity at each voxel, in units of percentage blood-oxygen level-dependent (%BOLD) change (Lindquist, 2008). Traditionally, task-activated brain regions are identified by conducting hypothesis tests on the %BOLD change for each voxel separately, adjusting for multiple testing (Lindquist, 2008).

While standard, the testing approach has two significant limitations. First, it is typically conducted under the null hypothesis that the change in brain activity is zero. However, in practice, a large amount of the brain may exhibit non-zero albeit low activation which may or may not be of interest (Gross and Binder, 2014). This means that increasing the sample size will result in rejecting the null in increasingly more locations, losing spatial precision (Bowring et al., 2019; Davenport et al., 2022). Instead, researchers may seek to identify brain regions where the activation is particularly strong, for example, greater than 2% BOLD change. Second, with hypothesis testing, fMRI results are typically presented with thresholded color-coded statistical maps that only highlight significant regions (Poldrack et al., 2008). However, test statistics are unitless and do not provide a clinical interpretation, prompting recommendations on more emphasis on effect estimates (Chen et al., 2017). Moreover, highlighting only significant areas overlooks areas that have large changes but are statistically insignificant due to insufficient power (Greenland et al., 2016). Instead, the problem of activation localization is more naturally formulated as finding confidence regions for the true activated region exceeding a threshold. This approach, analogous to presenting a confidence interval, allows non-zero thresholds, preserves information on the effect estimate and facilitates interpretation. Figure 1 illustrates a comparison between the traditional hypothesis testing approach and the confidence regions approach.

**Figure 1:**
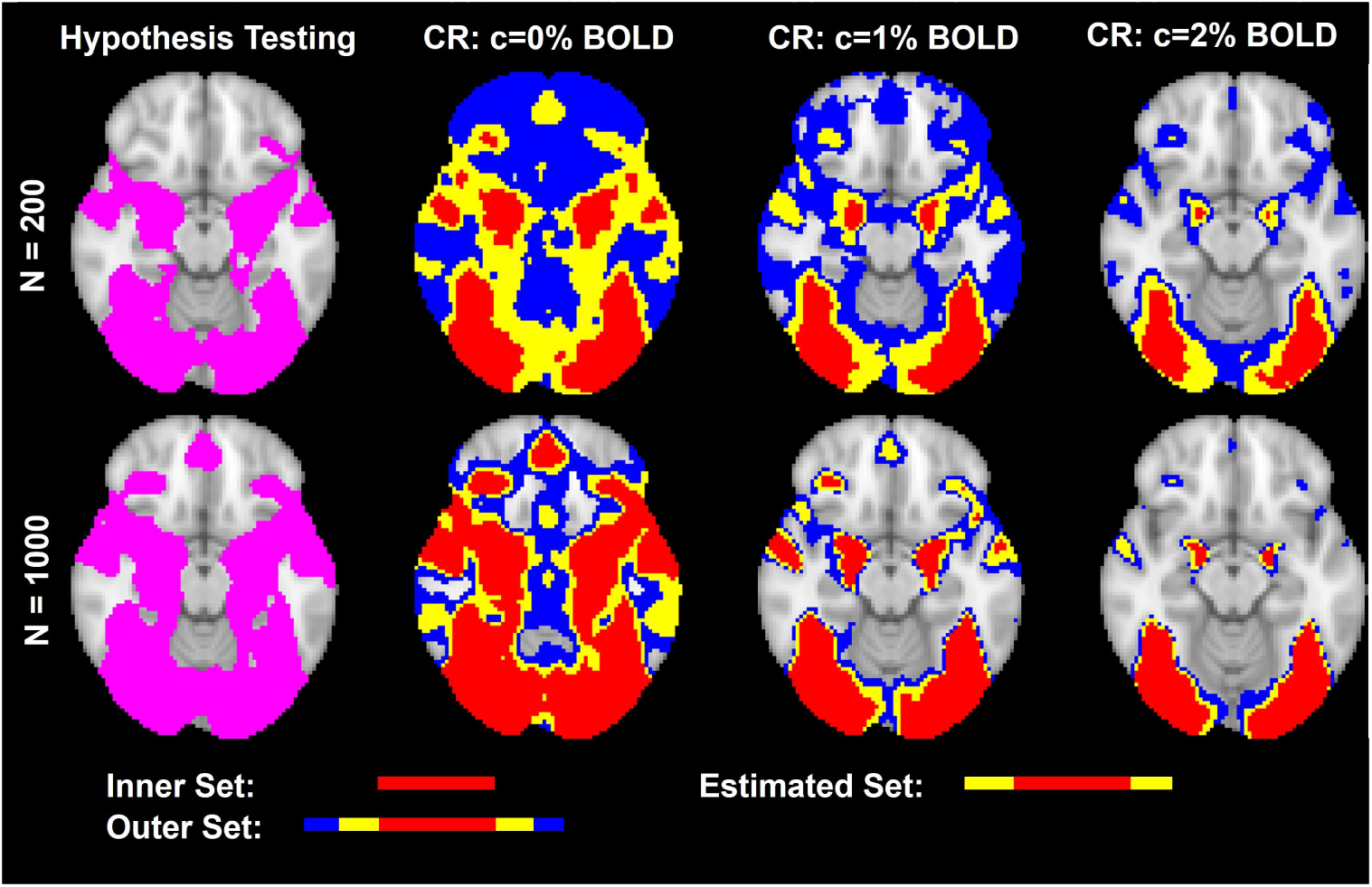
Activated brain regions obtained using classical hypothesis testing and the confidence regions (CR) approach with thresholds of 0, 1 and 2, with sample sizes of 200 and 1000. The data are from the Hariri faces/shapes “emotion” task in UK Biobank. Hypothesis testing was conducted using permutation based clusterwise inference at a cluster defining threshold of 3.1. For the CR results, the red region, union of red and yellow region, union of red and yellow and blue region represent the inner set, estimated set, outer set, respectively. To interpret the CR results, for example, at *c* = 2 % BOLD, we can state with at least 95% confidence that the true brain regions with more than 2% BOLD change lie between the inner set and the outer set. When sample size is large, hypothesis testing indicates many locations as statistically significant, losing spatial precision. In contrast, CRs using a non-zero threshold yield more informative and interpretable results.

Sommerfeld et al. (2018) proposed a spatial inference method for constructing confidence regions, which provide spatial uncertainty in the estimation of excursion sets of the mean function in images. This method was later refined and applied to fMRI data by Bowring et al. (2019), allowing inference on brain regions with non-zero activation thresholds. However, this general approach is limited to one predetermined activation threshold. In practice, deciding on a reasonable threshold beforehand may be difficult, and researchers are inclined to explore various thresholds, which necessitates addressing the issue of multiple testing over thresholds (Bowring et al., 2019). Moreover, this approach can only be applied to volume and not cortical surface data. This is a critical limitation since surface-based analyses, recognized for their greater sensitivity and reliability than volume-based methods, have received increasing attention (Tucholka et al., 2012). Bayesian approaches which provide posterior confidence regions for excursion sets of cortical surface data have been proposed (Mejia et al., 2019; Spencer et al., 2022). However, these also consider a single threshold and rely on assumptions of stationarity and Gaussianity.

Recently Ren et al. (2024) and Telschow et al. (2023) proposed a method for constructing confidence regions (CRs) that remain valid for all possible thresholds, hence the name, “simultaneous confidence regions (SCRs)”. In this method, CRs are produced by inverting simultaneous confidence bands (SCBs) at a certain threshold. The key step of this method is therefore the construction of valid SCBs, typically obtained via bootstrap techniques (Degras, 2011; Chernozhukov et al., 2013; Chang et al., 2017).

To extend the SCR method to fMRI studies, we need to ensure the validity of the bootstrap with fMRI data. Prior evaluations of the bootstrap have mostly used 1D or Gaussian models in simulations (Bowring et al., 2019; Telschow and Schwartzman, 2022), which fail to reflect the higher-dimensional, non-stationary, non-Gaussian nature of fMRI data (Hanson and Bly, 2001; Wager et al., 2005; Davenport et al., 2023). Eklund et al. (2016) emphasized that simulations under restrictive assumptions such as Gaussianity are insufficient to establish the validity of statistical methods in fMRI studies. They proposed using resting state validations, which fit a fake task design to resting state data in order to generate realistic noise and have become the gold standard for method validation in fMRI (Lohmann et al., 2018; Davenport et al., 2023; Andreella et al., 2023).

The contributions of this paper are as follows. First, we evaluate six bootstrap variants for constructing SCB, including the nonparametric bootstrap and multiplier bootstrap, through extensive 2D simulations with Gaussian and non-Gaussian data. We find that the Rademacher multiplier bootstrap-t performs the best, achieving a coverage rate close to the nominal level with sample sizes as low as 20. Second, we validate the corresponding coverage of the SCRs using realistic 3D resting-state fMRI data. Third, we have developed software that constructs confidence regions for data of any dimension, such as brain volume and surface data. Our software is equipped with visualization tools tailored for fMRI, including interactive apps that allow users to visualize activated brain regions as they adjust the activation threshold. Finally, we illustrate our approach with an application to both fMRI volume data and surface data from the Human Connectome Project.

We have implemented this method in the Python package SimuInf (Qin, 2024). A Matlab implementation is also available in the StatBrainz package (Davenport, 2024). Demonstrations of the interactive apps for volume and surface data analyses are provided in Figures 7 and 8. All the simulations and analyses were run on an Intel Core CPU@2.1 GHz with 16GB RAM.

## 2 Theory

### 2.1 Confidence Regions for an Excursion Set

Let *S* R*^D^*, *D ∈* N, be a domain (e.g. corresponding to the brain) and let *µ*: *S* → R be a signal of interest. The inverse image of *µ* under a set *U ⊂* R is defined as *µ^−^*^1^(*U*) = *{s* ∈ *S*: *µ*(*s*) *∈ U}*. For a real number *c*, if *U* = [*c,* ∞), then *µ*^-1^(*U*) is called the excursion set of *µ* above the level *c*. In the context of fMRI, researchers aim to identify areas of the brain activated during a task. Here *S ⊂* R^3^ corresponds to the set of voxels or vertices making up the brain and *µ*(*s*) represents the %BOLD change at voxel/vertex *s* ∈ *S*. For instance, setting *c* = 2, the excursion set *µ^−^*^1^[2, ∞), is the quantity of interest and represents brain areas with at least 2% BOLD change. CRs quantify the uncertainty in estimating *µ^−^*^1^[*c, ∞*). They consist of an inner set, denoted as CR_in_[c*, ∞*), and an outer set, denoted as CR_out_[c*, ∞*), such that

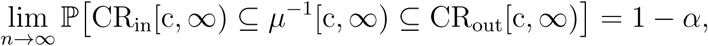

where *α* is the Type 1 error rate, typically set at 0.05. Of note, the inner and outer sets are estimated from data, making them random quantities. While Bowring et al. (2019) refers to them as upper and lower sets respectively, we prefer the terms “inner” and “outer” to indicate that the inner set is contained within the outer set. Moreover, Ren et al. (2024) used the term “confidence sets”; however, we favor the term “confidence regions” as it emphasizes that they quantify spatial uncertainty.

### 2.2 Constructing Simultaneous Confidence Regions by Inverting the SCB

To obtain SCRs suitable for application in brain imaging, we follow the approach of Ren et al. (2024). They proposed constructing CRs of *µ^−^*^1^[*c,* ∞) that are valid for all *c* R by inverting a SCB of *µ*(*s*). An asymptotic SCB consists of a lower function 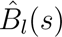 and an upper function 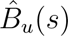 such that:

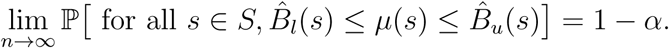

Given an asymptotic SCB, CRs can be calculated as 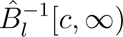 for the inner set and 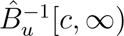 for the outer set. Theorem 1 in Ren et al. (2024) established an equivalence between the SCB and the CRs, that is:

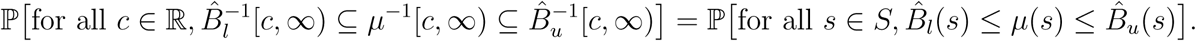

These CRs are valid for all *c ∈* R, hence the name, “simultaneous confidence regions”. That is, we have:

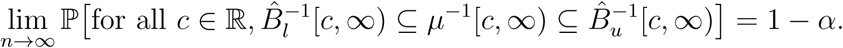

Figure 2 (A) illustrates the idea of this method with a 1D function *µ*(*s*): *s ∈ S ⊂* R. To estimate the excursion set *µ^−^*^1^[*c,* ∞), we first calculate 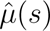, the estimator of *µ*(*s*). The SCB of *µ*(*s*) is then constructed, consisting of 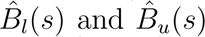. Finally, the inner, estimated, and outer sets are obtained by inverting 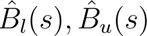 respectively at the threshold *c*. With a 2D function, as depicted in Figure 2 (B), the estimated set and its SCRs can be obtained similarly.

**Figure 2:**
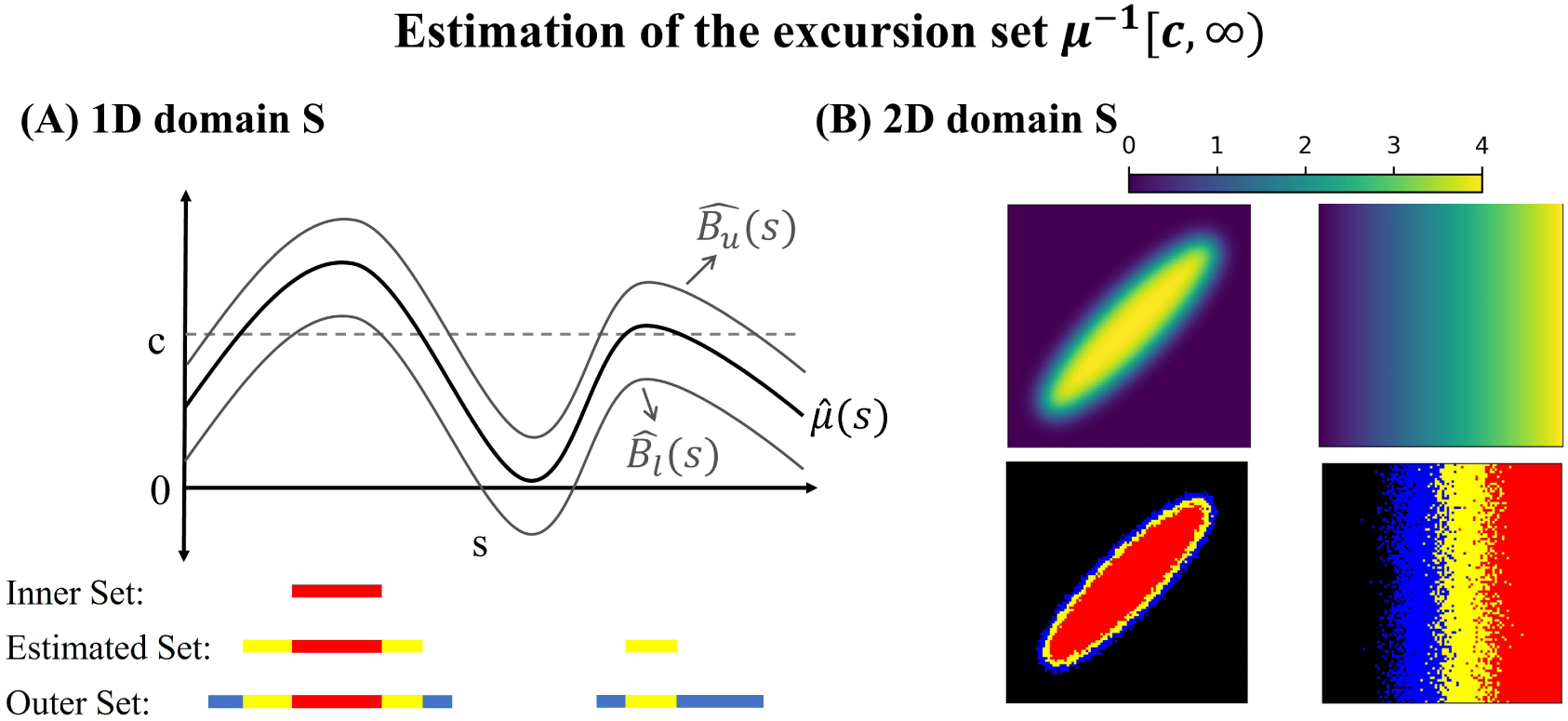
Illustration of the simultaneous confidence regions method with a 1D function (A) and a 2D function (B). The red region, union of red and yellow region, union of red, yellow and blue region represent the inner set, estimated set, and outer set, respectively. In (A), the black curve represents the estimator of *µ*(*s*). The two gray curves represent the simultaneous confidence band of *µ*(*s*). In (B), the top two panels represent two examples of *µ*(*s*), taking a shape of an ellipse and a ramp. The bottom two panels represent their corresponding estimated excursion sets and confidence regions based on 40 samples from model 1.

### 2.3 SCB in Functional Signal-plus-noise Models

This study focuses on signal-plus-noise models, which include regression models that are widely used in second-level fMRI data analyses (Mumford and Nichols, 2009). Let 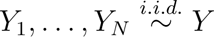 be an independent and identically distributed (i.i.d) sample of random functions, where *Y* follows the following functional signal-plus-noise model:

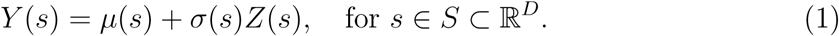

Here, *µ*(*s*) and *σ*(*s*) are fixed functions, *Z*(*s*) is a random function with mean zero and variance one for all *s*, *ɛ*(*s*) = *σ*(*s*)*Z*(*s*) is the noise function. Of note, we do not assume stationarity, a particular correlation structure or a particular distribution (for example, Gaussian) on the noise field *ɛ*(*s*).

Define the sample mean and sample variance as:

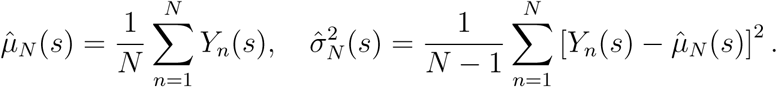

Of note, the subscript *N* in 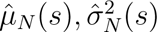 emphasizes that these estimators depend on the sample size *N*. An asymptotically valid Wald based SCB of *µ*(*s*) is:

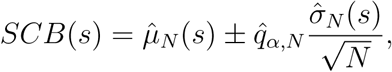

where the quantile 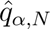 can be obtained from bootstrap methods as described in Section 2.4.

### 2.4 Variants of Bootstrap Methods

SCBs are typically constructed using the bootstrap. In this section we describe how two of the most widely used bootstrap methods can be used to provide the quantile 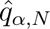 and summarize additional variations at the end.

Nonparametric bootstrap(Degras, 2011):

1. Resample from *Y*_1_*,…, Y_N_* with replacement to produce a bootstrap sample 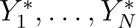
2. Compute 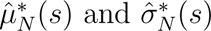 using the sample 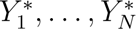.
3. Compute 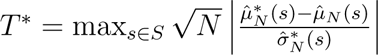

4. Repeat steps 1 to 3 many times to get the distribution of *T*\* and set 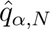 to be the (1 *− α*)*th* quantile of this distribution.

Multiplier (or Wild) Bootstrap (Chang et al., 2017):

1. Define residuals 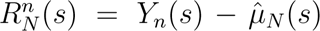, compute 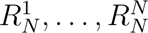 and multipliers 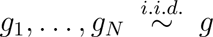 with *E*[*g*] = 0 and var[*g*] = 1 to produce a bootstrap sample 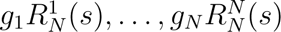. Common choices of *g* are a standard Gaussian random variable or a Rademacher random variable, which takes values of 1 and −1 with probability 1*/*2.
2. Compute 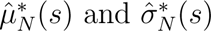 from 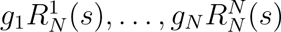.
3. Compute 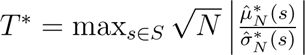
4. Repeat steps 1 to 3 many times to get the distribution of *T*\* and set 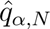 to be the (1 *− α*)*th* quantile of this distribution.

Of note, in both methods described above, the third step standardizes the bootstrap sample mean with bootstrap sample standard deviation (SD), akin to the calculation of a *T* score. An alternative approach is to standardize with the original sample SD, mirroring the calculation of a *Z* score (Chernozhukov et al., 2013; Sommerfeld et al., 2018). These two types of standardizations are referred to as *T* and *Z* standardization.

## 3 Methods

### 3.1 2D Simulations

We conducted a series of 2D simulations to evaluate various bootstrap methods for constructing SCBs, assessing the following aspects: coverage rate, runtime, precision and stability. We considered various scenarios and bootstrap methods, as detailed below. For all scenarios considered, the number of simulation replications was 1000, the number of bootstrap samples was 1000 and the significance level *α* was 0.05, corresponding to a target coverage level of 1 − *α* = 0.95. Coverage rate was calculated as the proportion of simulation instances in which the true means at all grid points fell within their respective confidence bands, thereby assessing the simultaneous coverage across all grid points. Average runtime across the 1000 simulation replications was calculated. Precision was assessed by the mean of the quantile 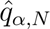 across the 1000 simulation replications, where a smaller value corresponds to a narrower and thus more precise SCB. Stability was assessed by the standard deviation (SD) of 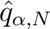 across the 1000 replications, where a smaller value represents a more stable SCB.

In each simulation instance, the data were generated as an i.i.d sample from model 1. The following parameters were varied, leading to a combination of 400 scenarios:

- shape of the signal *µ*(*s*) *∈ {*ellipse, ramp*}*, as depicted in Figure 2(B)
- noise distribution before smoothing *∈ {*Standard Gaussian, Student’s *t* with 3 degrees of freedom (*t*_3_)}

In detail, before smoothing, the *ɛ*(*s*) was generated as i.i.d over *s* from the given distribution. The *t*_3_ distribution was chosen since it approximates the noise distribution of fMRI data (Davenport et al., 2023)

- full width at half maximum (FWHM) in Gaussian kernel smoothing of the noise ∈ {0, 1, 2, 3, 4}

Of note, smoothing introduces the correlation in the noise *ɛ*(*s*) over *s*.

- SD of the noise after smoothing ∈ {1, 10}

Specifically, after smoothing, the noise *ɛ*(*s*) was normalized to have the same SD of 1 or 10 over *s*.

- 2D image size ∈ {50 *×* 50, 100 *×* 100}
- sample size ∈ {20, 40, 60, 80, 100}

For each scenario, we evaluated six bootstrap methods, which are a combination of three bootstrap types (nonparametric, Gaussian multiplier, Rademacher multiplier) and two standardization types (*T*, *Z*).

### 3.2 3D Validations

In order to test the performance of the SCRs in realistic noise settings, we conducted resting state validations to assess the coverage rate of the SCBs and the resulting confidence regions. To do so we used 3D contrast images obtained from resting-state fMRI data of 198 healthy controls (Beijing dataset) from the 1,000 Functional Connectomes Project (Biswal et al., 2010). These images were processed using FSL (Jenkinson et al., 2012) by Eklund et al. (2016) using a fake task design consisting of a 10-s on/off block activity paradigm and a 4mm FWHM smoothing. Since resting-state data should not contain systematic changes in brain activity, these contrast images are expected to have a mean of zero. A realistic signal was introduced by adding the average %BOLD change during the Hariri faces/shapes “emotion” task, from 4,000 UK Biobank participants (Alfaro-Almagro et al., 2018), to each image.

To evaluate the coverage rate for a sample of size *n*, in each analysis instance, *n* images were sampled without replacement from the 198 3D contrast images. SCBs were subsequently constructed using the Rademacher multiplier bootstrap-t and the confidence regions for various numbers of predefined thresholds were obtained. The Rademacher multiplier bootstrap-t was used since it achieved a coverage rate close to the nominal level in previous 2D simulations. This procedure was replicated 1,000 times, mimicking the regular Monte Carlo simulation but with realistic datasets. The coverage rate of the SCBs was calculated as described in Section 3.1. The coverage rate of the confidence regions was calculated as the proportion of analysis instances in which the true excursion set contained the inner set and was contained by the outer set for all predefined thresholds, thereby assessing the simultaneous coverage across thresholds. That is,

SCR coverage rate = #*{*Analysis Instance: for all *c ∈ K,* CR_in_ *⊆ µ^−^*^1^[c*, ∞*) *⊆* CR_out_*}/*100 where *K* is the set of predefined thresholds.

The thresholds considered were taken to be equidistant from −20 to 20, covering the range where the majority of the signal lies in. Different sample sizes (10, 20, 30, 40, 50) and numbers of thresholds (5, 10, 50, 100, 1000) were examined to evaluate the method’s performance under different scenarios. Since the assumed activity paradigm in the first-level analysis may influence the results (Eklund et al., 2016), the above evaluations were repeated with contrast images generated with an event activity paradigm (1-to 4-s activation, 3-to 6-s rest, randomized), 4mm FWHM using FSL.

### 3.3 Application to Task fMRI Volume and Surface Data

To illustrate the performance of the SCRs in practice, we applied them to volume and cortical surface task fMRI data from the Human Connectome Project (HCP). The sample included 78 unrelated subjects engaged in a working memory task. A second-level analysis was conducted on the 78 3D contrast images to determine the task-activated brain regions across the participants. A similar analysis was conducted for the 78 cortical surface images to determine activated surface areas. Detailed descriptions of the study protocol, task paradigm and first-level analyses are available in Barch et al. (2013) and Glasser et al. (2013), with a brief summary provided below.

The task contained two runs, each consisting of four blocks. In each block, the participant undertook either a 2-back memory task or a 0-back control task. The experimental design was arranged such that, in each run, two blocks were designated to the 2-back memory task, and two blocks were designated to the 0-back control task. In each block, a participant was shown a stimuli image (a picture of a face or a place, for instance) and then asked to recall the image they were shown. They were either asked to recall the most recent image (the 0-back image) or the image shown to them two images prior (the 2-back image). First-level analyses were conducted independently for each participant using FSL, where the task design was regressed onto BOLD response, generating a contrast image for each participant. These images represent the difference in BOLD response between the 2-back task and the 0-back task.

## 4 Results

### 4.1 2D Simulations

The simulation results are presented for the scenarios with an ellipse shape, FWHM smoothing of 2 and an image size of 100 × 100. Results in other scenarios are similar and are provided in the supplementary file. In assessing the coverage rate, as depicted in Figure 3(A), among all the methods evaluated, the Rademacher multiplier bootstrap-t performs the best. It maintains a coverage rate consistent with the nominal level of 0.95 across variations in sample size, noise distribution before smoothing, and noise SD after smoothing. In general, methods with *T* standardization have a coverage rate closer to the nominal level than their counterparts with *Z* standardization, especially when sample size is small.

**Figure 3:**
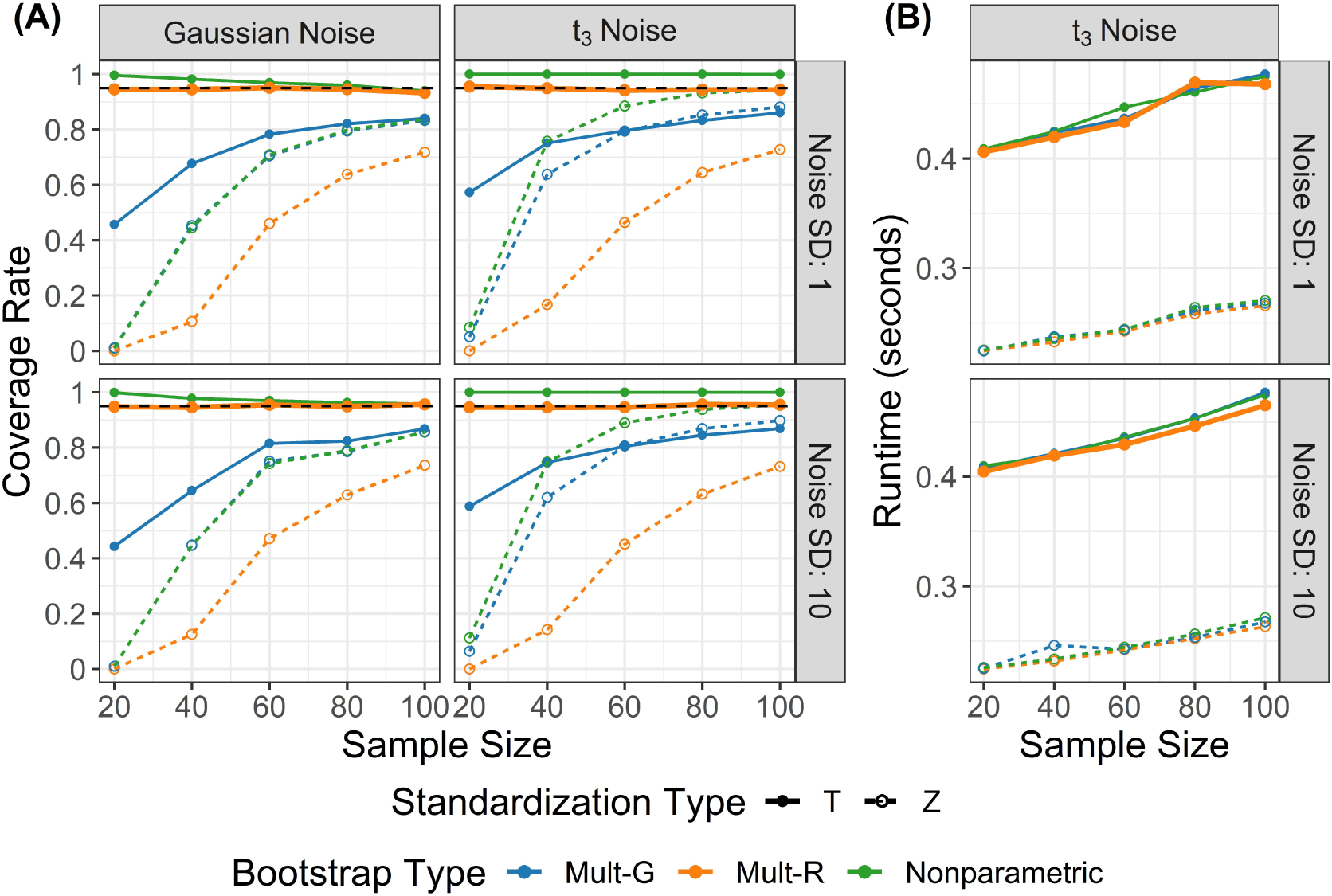
Results of 2D simulations on coverage rate (A) and runtime (B) under variations in sample size, noise distribution before smoothing and noise standard deviation (SD) after smoothing. The black dashed line represents the target coverage rate of 0.95. Six bootstrap methods (3 bootstrap types × 2 standardization types) were evaluated. (A) Among these methods, the Rademacher multiplier bootstrap-t performs the best, achieving a coverage rate close to the target level under all variations considered. (B) Methods with *Z* standardization are faster than *T* standardization, independent of bootstrap type. Runtime results under Gaussian noise are very similar and can be found in the supplementary file.

When the noise follows Gaussian distribution, the nonparametric bootstrap-t method is overly conservative with small samples, yet aligns more with the nominal level as sample size increases. Conversely, when the noise follows a *t* distribution with 3 degrees of freedom (*t*_3_), the nonparametric bootstrap-t remains excessively conservative and shows no improvement with larger samples.

Regarding runtime, as illustrated in Figure 3(B), methods with *Z* standardization are faster across all sample sizes. They complete in less than 0.3 seconds for a single simulation instance involving 1000 bootstrap iterations, which is approximately half the runtime required by *T* standardization. Within the same standardization, the three types of bootstrap methods have very similar runtime.

Figure 4 presents the results for the mean and SD of estimated SCB quantiles, assessing precision and stability of each method. Only the two methods achieving coverage rates close to the nominal level are shown, since it is meaningless to consider precision and stability for methods with poor coverage. Under all scenarios, the Rademacher multiplier bootstrap-t gives a more precise and stable SCB than its main competitor.

**Figure 4:**
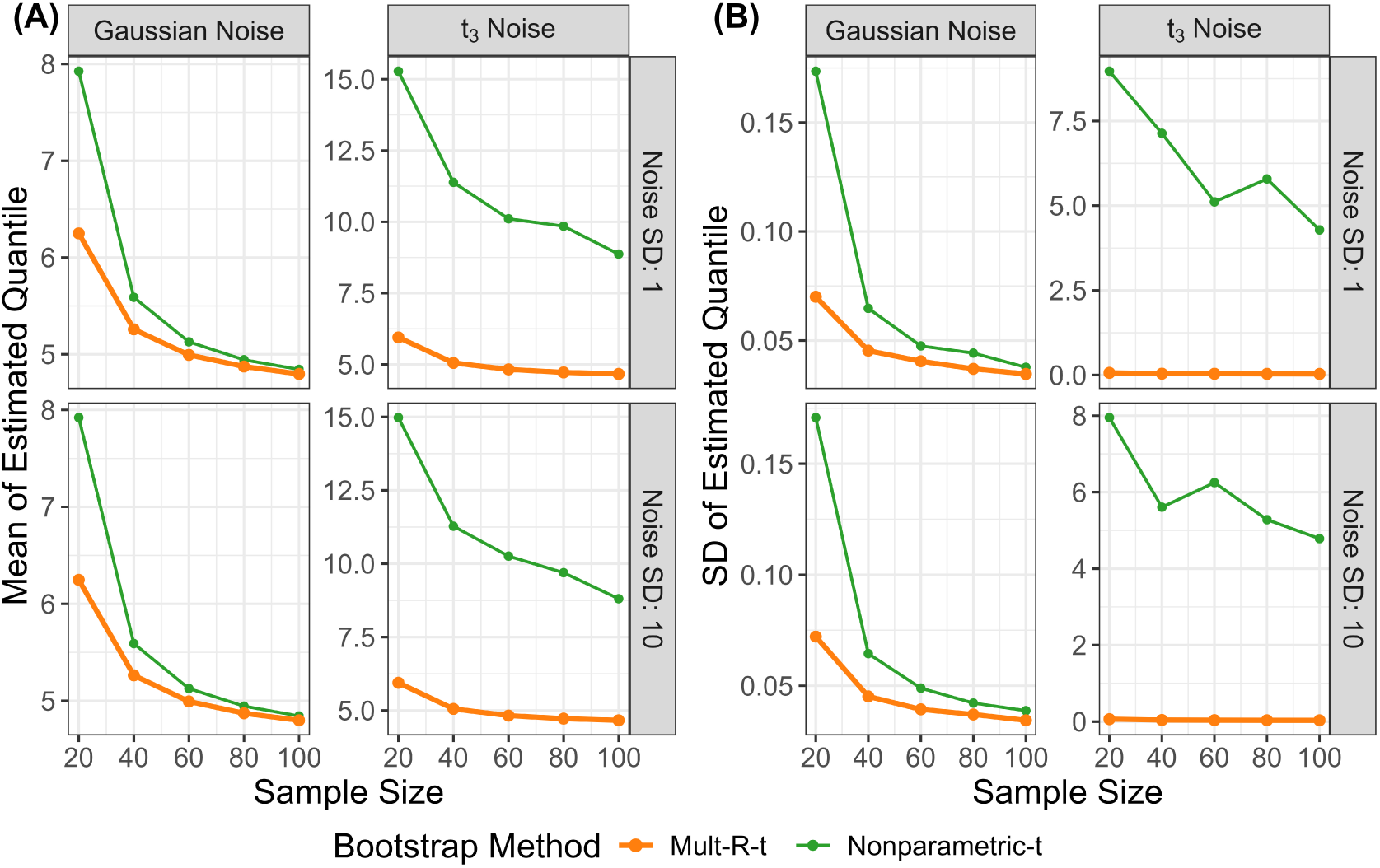
Results of 2D simulations on mean (A) and SD (B) of estimated SCB quantiles, under variations in sample size, noise distribution and noise SD. Two bootstrap methods that achieved a good coverage rate were compared. A smaller mean quantile represents a narrower (i.e., more precise) SCB and a smaller SD of quantiles represents a more stable SCB. The Rademacher multiplier bootstrap-t gives a more precise and stable SCB than its competitor, the nonparametric-t under all scenarios.

### 4.2 3D Validations

We conducted 3D validations using the SCR method with the Rademacher multiplier bootstrap-t, which achieved the target SCB coverage rate in previous 2D simulations. As depicted in Figure 5, the coverage rates of the SCBs closely align with the nominal level of 0.95, independent of the sample size and assumed activity paradigm, validating the use of the Rademacher multiplier bootstrap-t for SCB construction in realistic fMRI data. Regarding the resulting confidence regions, their coverage rates approach from above to the nominal level as the number of considered threshold levels increases.

**Figure 5:**
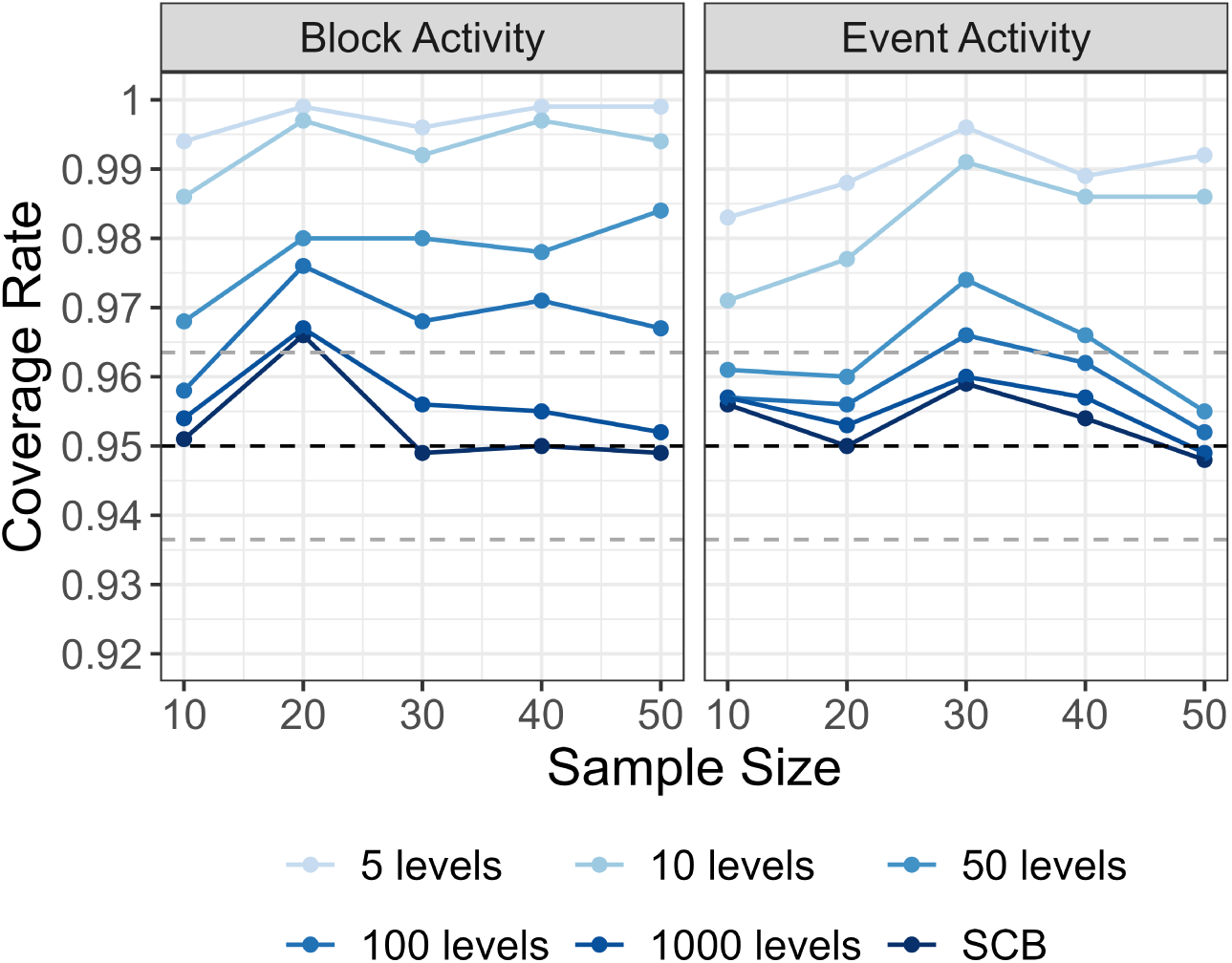
Coverage rate results of 3D validations with realistic fMRI data under variations in sample size and assumed activity paradigm. The confidence regions were constructed by inverting the SCBs obtained by the Rademacher multiplier bootstrap-t. The black dashed line represents the target coverage rate of 0.95. The two gray dashed lines capture the uncertainty due to simulation and correspond to 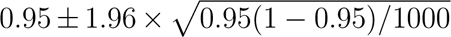. The coverage rate of the SCB is close to the target level. The coverage rate of the confidence regions approaches from above to the nominal level as the number of threshold levels increases.

### 4.3 Application to Task fMRI Volume and Surface Data

The SCR method with the Rademacher multiplier bootstrap-t was applied to both the fMRI volume and surface data from HCP, collected during a working memory task. The results are presented in Figure 6(A) for volume data and Figure 6(B) for surface data. Demonstrations of interactive apps to visualize the results as users adjust the activation thresholds are provided in Figures 7 and 8. Results with additional thresholds and slices in different directions are provided in the supplementary file.

**Figure 6:**
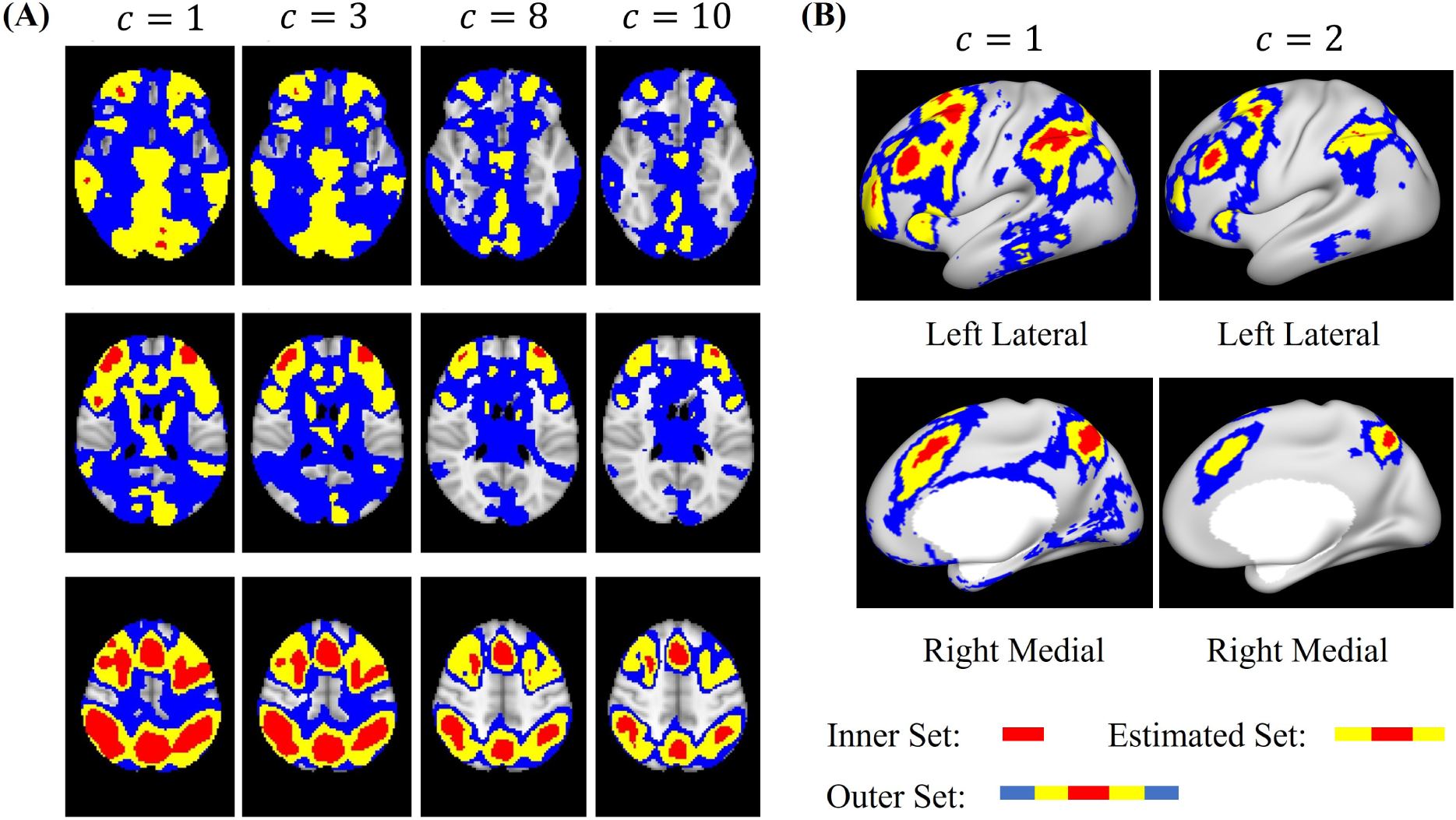
Confidence region results of fMRI volume data (A) and surface data (B) obtained during a working memory task. The red region, union of red and yellow region, union of red and yellow and blue region represent the inner set, estimated set, outer set, respectively. Each column displays the results for a particular threshold *c* by showing three distinct slices of the 3D brain in (A) or by showing the left and right hemispheres in (B). For example, the second column panel of (A) shows the result for brain regions with at least 3% BOLD change.

In both analyses, the activation thresholds selected for presentation were those that yielded the most informative and interesting results after exploring a range of thresholds. A major advantage of this method is its capacity to provide valid inference at all potential thresholds, offering great flexibility. For example, with the second column in Figure 6(A), we can conclude with at least 95% confidence that the brain region within the red area has an activation of at least 3% BOLD change. Similar conclusions can be made for all the other thresholds considered. Interactive apps to visualize the results as users adjust the activation threshold are demonstrated in Figures 7 and 8.

**Figure 7:**
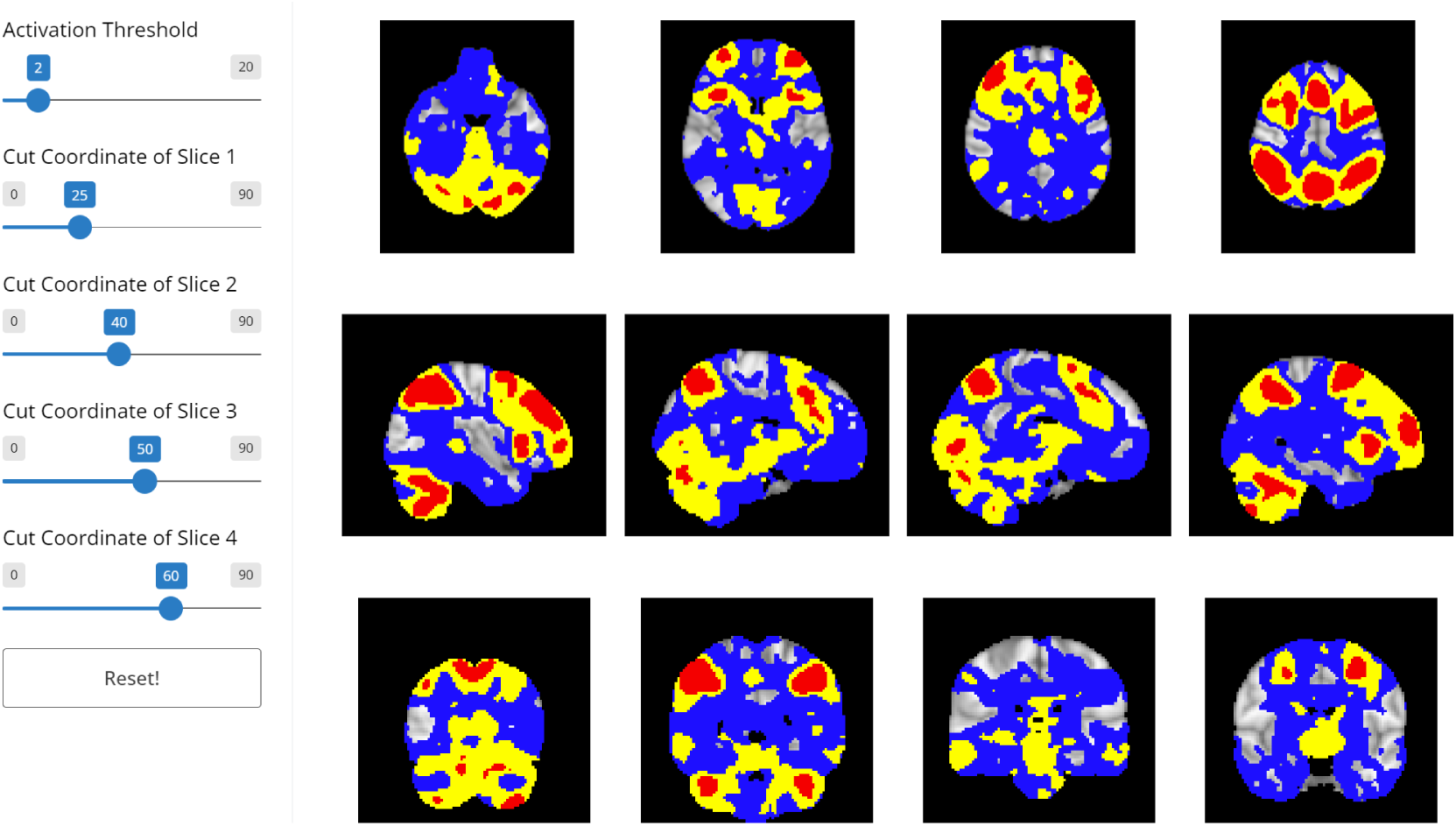
A demonstration of the interactive visualization tool for volume data analysis. This tool allows users to view the results of the confidence regions and estimated excursion sets as they change the activation threshold and the coordinates of four slices. Each column corresponds to a particular slice at a given coordinate. Each row corresponds to a particular direction of the slice: axial, sagittal and coronal, listed from top to bottom.

**Figure 8:**
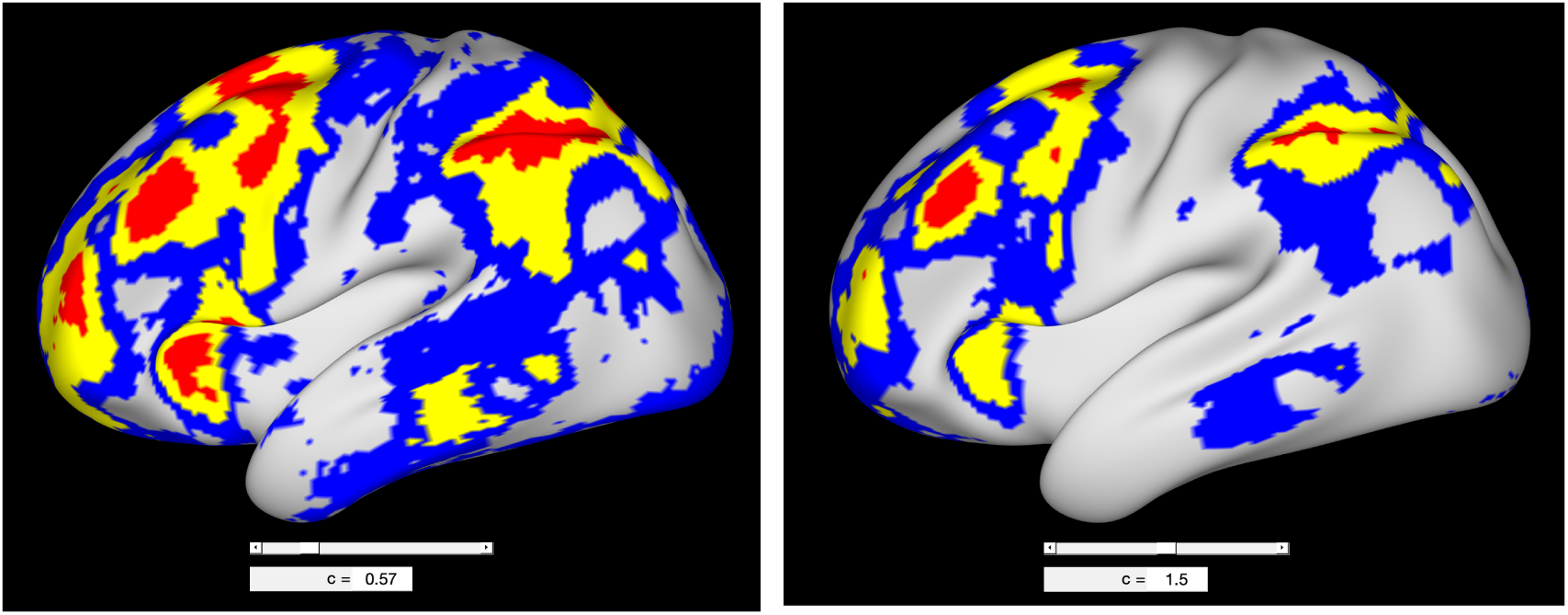
A demonstration of the interactive visualization tool for surface data analysis. This tool allows users to view the results of the confidence regions and estimated excursion sets as they change the activation threshold. In the example two thresholds, *c* = 0.57 and 1.5 are shown (which are in units of % BOLD change).

## 5 Discussion

In this study, we extended the SCR method in Ren et al. (2024) to the neuroimaging setting. We evaluated six bootstrap approaches for SCB construction using 2D simulations. The Rademacher multiplier bootstrap with *T* standardization performed the best, achieving a coverage rate close to the nominal level with sample sizes as low as 20. We further validated this method using real resting-state 3D fMRI data, a technique that has become the gold standard, by creating realistic noise that reflects the non-Gaussian and non-stationary structure of fMRI data. Our applications to real task fMRI volume value and surface data showcase the utility of this method in neuroimaging. Moreover we have developed software packages which implement this method and are equipped with visualization tools designed for fMRI. In conclusion, we confirm the validity of this method with the Rademacher multiplier bootstrap-t and advocate for its broader application in fMRI studies for localizing activated brain regions.

A key advantage of SCRs is that they provide valid inference simultaneously across all activation thresholds. This enables researchers to fully explore the data and choose the thresholds which provide the most interesting results, without concerns about multiple comparison issues over thresholds. We have developed interactive tools for both volume and surface data analyses, allowing users to visualize the activated brain regions as they adjust the threshold. Another strength of our method is that it does not require stationarity, a particular correlation structure or distribution on the noise field. This reduces bias from model misspecification compared to other methods such as classical implementations based on random field theory (Worsley et al., 1996, 2004), which have been shown to perform poorly in fMRI due to the non-stationarity and high levels of non-Gaussianity (Eklund et al., 2016).

Our 2D simulations assessed six bootstrap methods on coverage rate and runtime. Regarding coverage rate, the superior performance of the Rademacher multiplier bootstrap-t aligns with previous studies which considered simpler 1D or Gaussian scenarios (Telschow and Schwartzman, 2022; Bowring et al., 2019). Regarding runtime, we found a longer run-time of bootstrap approaches with *T* standardization than *Z* standardization, regardless of the bootstrap type. This is expected since *Z* standardization only uses the SD of the original sample whereas *T* standardization requires calculating the SD of each bootstrap sample. Nonetheless, with a 100 × 100 image, methods with *T* standardization completed within 0.6 seconds on a regular laptop, suggesting runtime concerns are minimal. Considering both aspects of coverage rate and runtime, we recommend the use of the Rademacher multiplier bootstrap-t.

Our 3D resting-state validations showed that SCRs using the Rademacher multiplier bootstrap-t controls the coverage rate at or above the nominal level in realistic fMRI data. As the number of thresholds increases, the coverage rate of the confidence regions approaches that of the SCBs from above. This occurs because the probability of coverage at a finite number of thresholds is always greater than for all thresholds, with equality in the limit. This allows the user to choose the threshold, even data driven, without worrying about incurring additional error.

Using our approach we explored the brain regions which are activated during a working memory task in both volume and surface data. The results are in line with previous research that associates working memory with fronto-parietal brain regions (Engström et al., 2015; Chai et al., 2018). However, prior results were obtained using hypothesis testing under the null of no activation, without providing spatial uncertainty (see for example, Figure 3 in Engström et al. (2015)). In contrast, our method shows the spatial uncertainty and captures the strength of the activation in interpretable units of %BOLD change.

Our work can be extended in the following directions. First, our implementation of the method focuses on a second-level analysis to estimate population mean, where it is reasonable to assume the contrast images from different individuals are i.i.d. In first-level analyses, where the time series of the BOLD response during an fMRI experiment is analyzed, the i.i.d assumption is violated. In such cases, SCB construction methods tailored for time series, for instance using the block bootstrap (Politis, 2003) to estimate the quantile, could be used. Once a valid SCB is established, SCRs can be constructed similarly by inverting the SCB. Second, non-bootstrap methods for constructing SCB could be considered, such as those based on the functional central limit theorem (Degras, 2011) or the Gaussian kinematic formula (Telschow and Schwartzman, 2022; Telschow and Davenport, 2023). Third, extensions to non-linear test statistics could also be considered, which could be obtained by bootstrapping delta residuals (Telschow et al., 2022). Finally, since our method enjoys valid inference for all thresholds simultaneously, it is conservative when users have specific pre-determined thresholds of interest. While uncommon, in that case, we recommend using the method in Bowring et al. (2019) for a single threshold and the method in Telschow et al. (2023) for a range of thresholds to achieve greater spatial precision.

## Data and Code Availability

Data and code are available at https://github.com/JiyueQin/SimuInf. The Human Connectome Project data can be provided upon request after users sign the data use agreement required by HCP, as instructed in the ReadMe file of the above GitHub link.

## Author Contributions

J.Q. drafted the manuscript, implemented the method in Python, conducted simulations and data analysis. S.D. implemented the method in MATLAB and contributed to the simulations and data analysis. A.S. and S.D. conceived the idea and oversaw the project. All authors edited and revised the manuscript.

## Acknowledgments

S.D. and A.S. were partially supported by NIH grant R01MH128923. Part of this research has been conducted using the UK Biobank Resource under Application Number: 34077. Data were provided in part by the Human Connectome Project, WU-Minn Consortium (Principal Investigators: David Van Essen and Kamil Ugurbil;1U54MH091657) funded by the 16 NIH Institutes and Centers that support the NIH Blueprint for Neuroscience Research; and by the McDonnell Center for Systems Neuroscience at Washington University.

## Ethics Statement

This study adheres to ethical guidelines provided by the Committee on Publication Ethics (COPE) and International Committee of Medical Journal Editors (ICMJE). Our study involves only the analysis of public data and thus is exempt from IRB review.

## Supplementary Results

## 1 Results of Task fMRI Volume Data Analysis

**1.1 Axial Slices**

**Figure 1:**
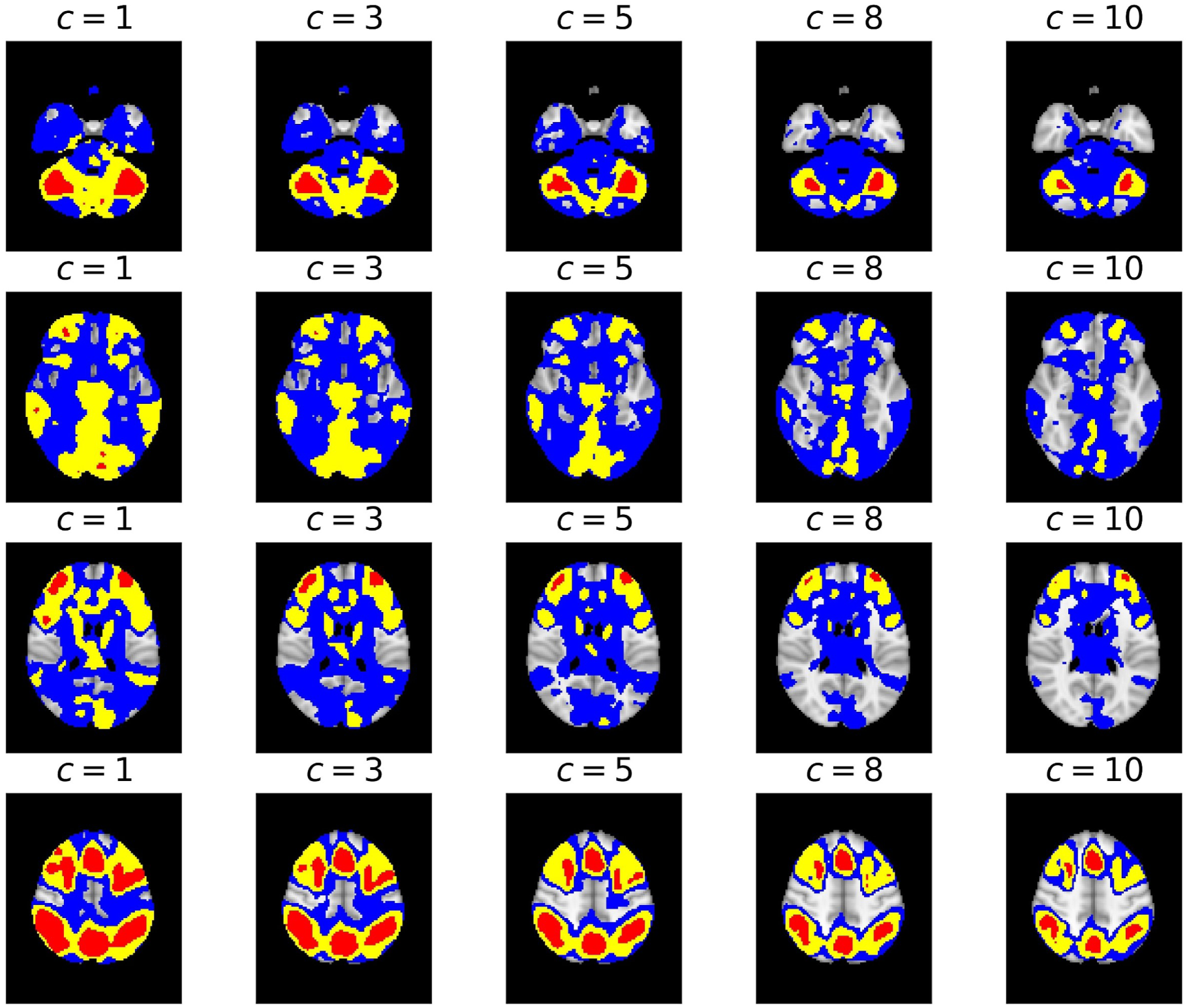
Confidence region results of fMRI volume data obtained, displayed in axial slices. The red region, union of red and yellow region, union of red and yellow and blue region represent the inner set, estimated set, outer set, respectively. Each column represents a particular activation threshold *c* and each row represents a particular axial slice.

**1.2 Sagittal Slices**

**Figure 2:**
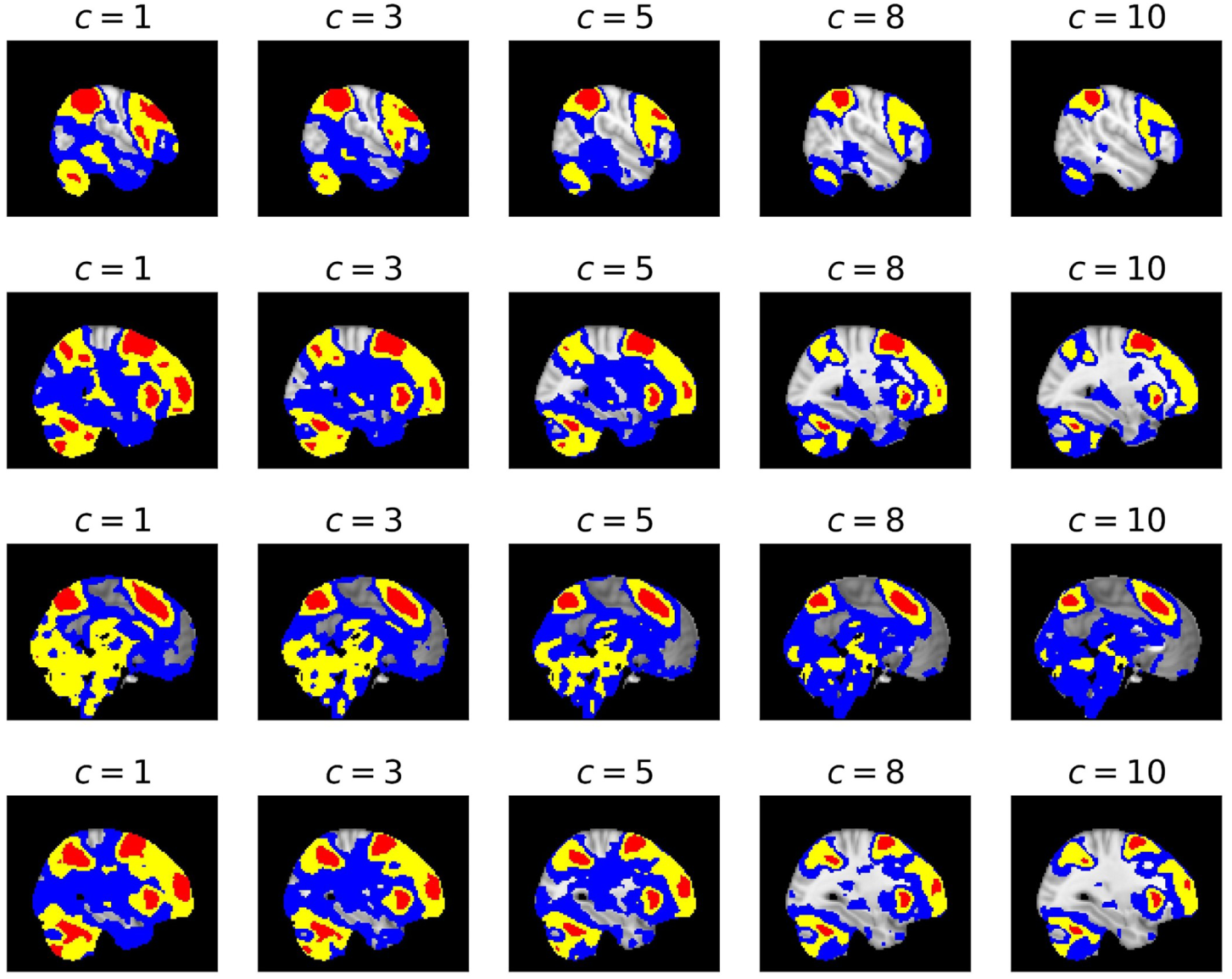
Confidence region results of fMRI volume data, displayed in sagittal slices. The red region, union of red and yellow region, union of red and yellow and blue region represent the inner set, estimated set, outer set, respectively. Each column represents a particular activation threshold *c* and each row represents a particular sagittal slice.

**1.3 Coronal Slices**

**Figure 3:**
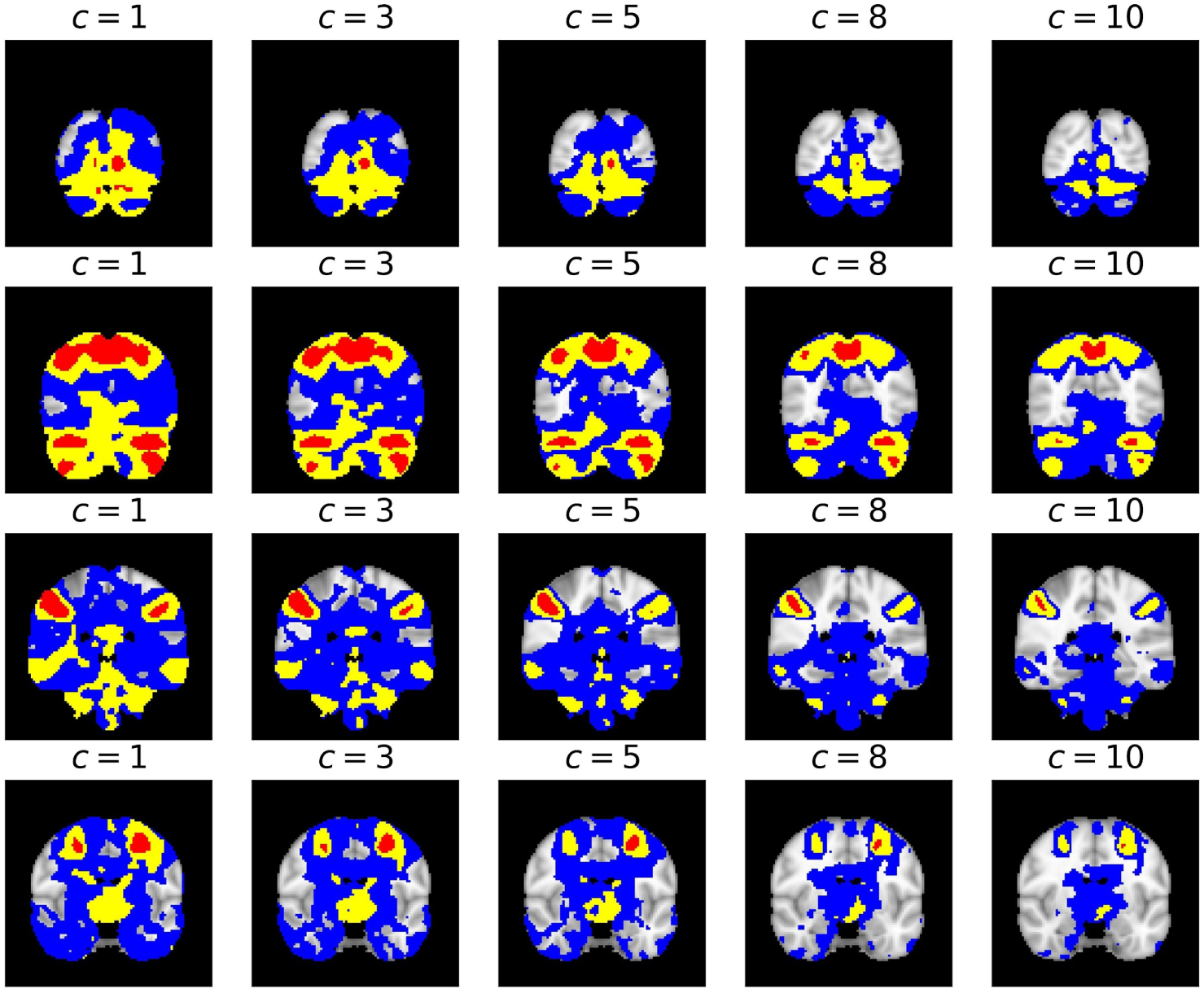
Confidence region results of fMRI volume data, displayed in coronal slices. The red region, union of red and yellow region, union of red and yellow and blue region represent the inner set, estimated set, outer set, respectively. Each column represents a particular activation threshold *c* and each row represents a particular coronal slice.

## 2 2D Simulation Results

**2.1 Coverage Rate**

Results of coverage rate in all simulated scenarios:

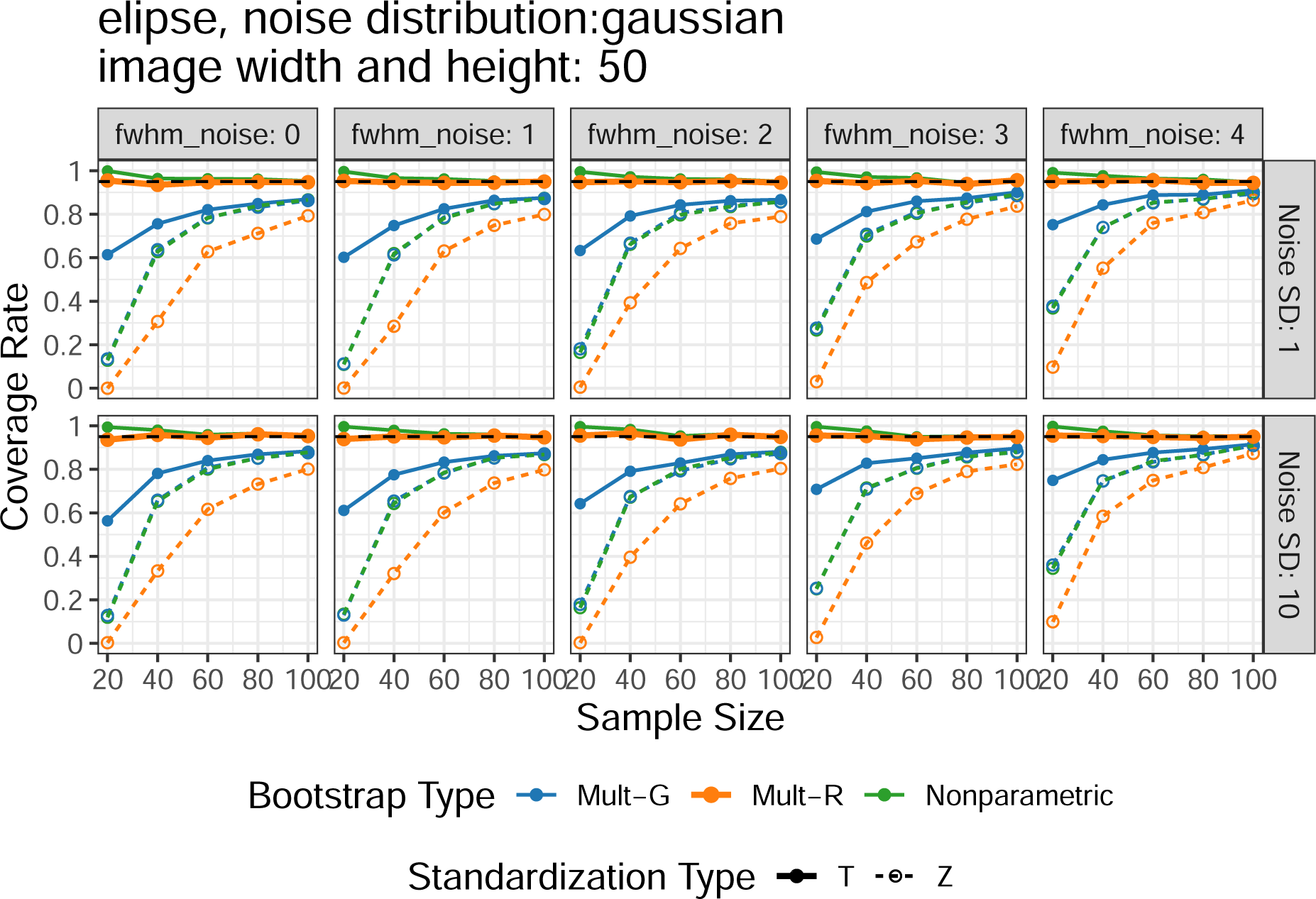

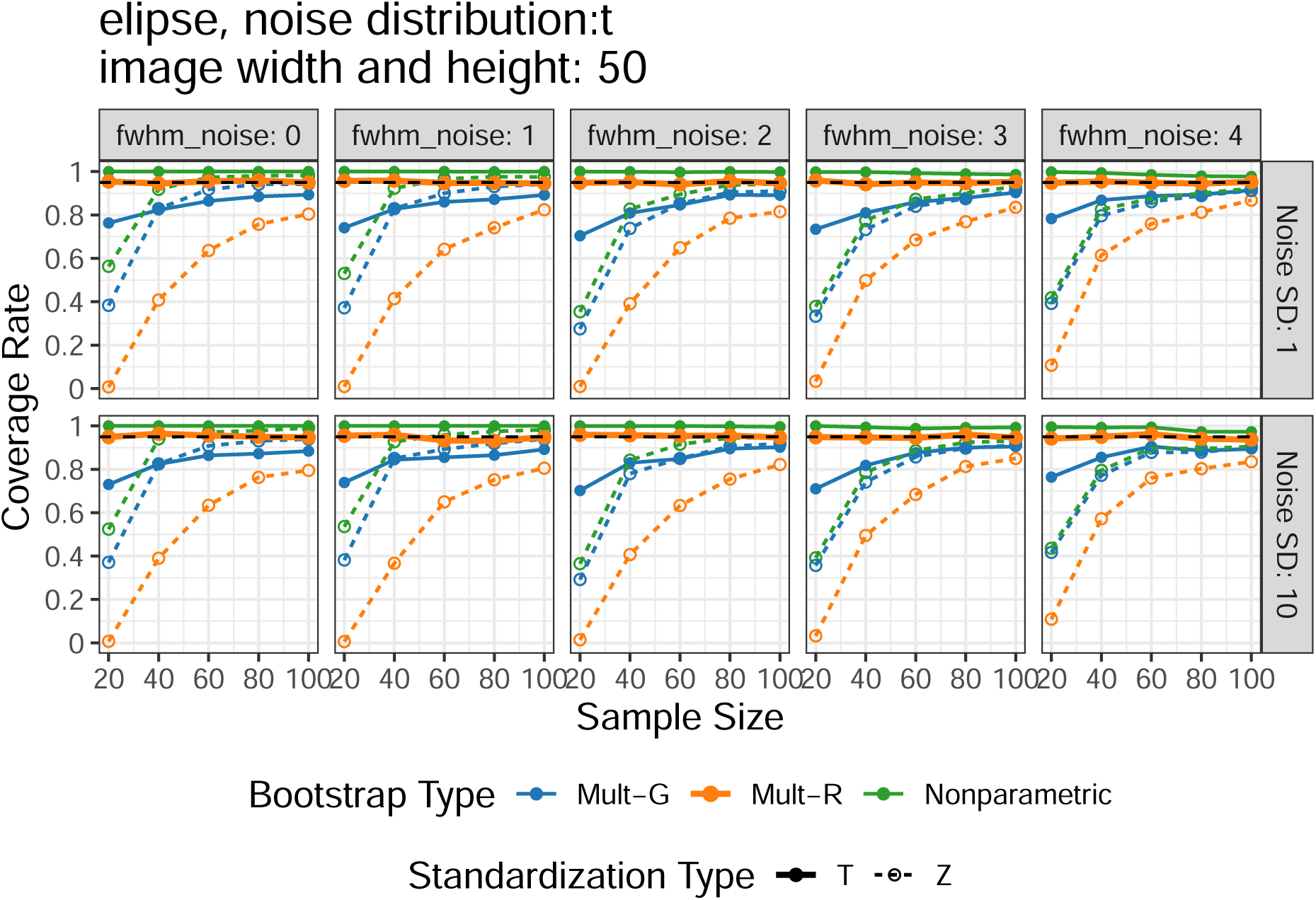

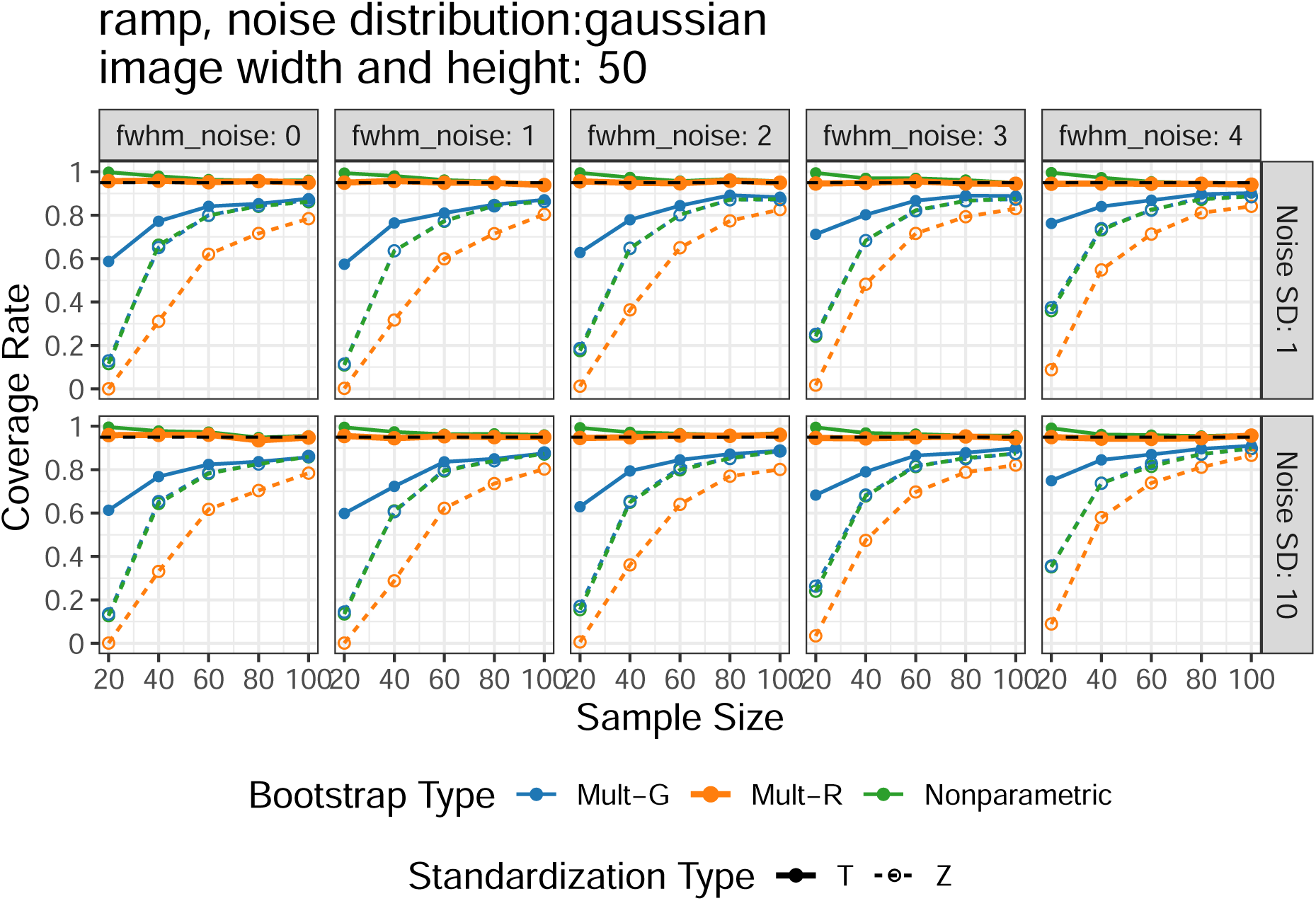

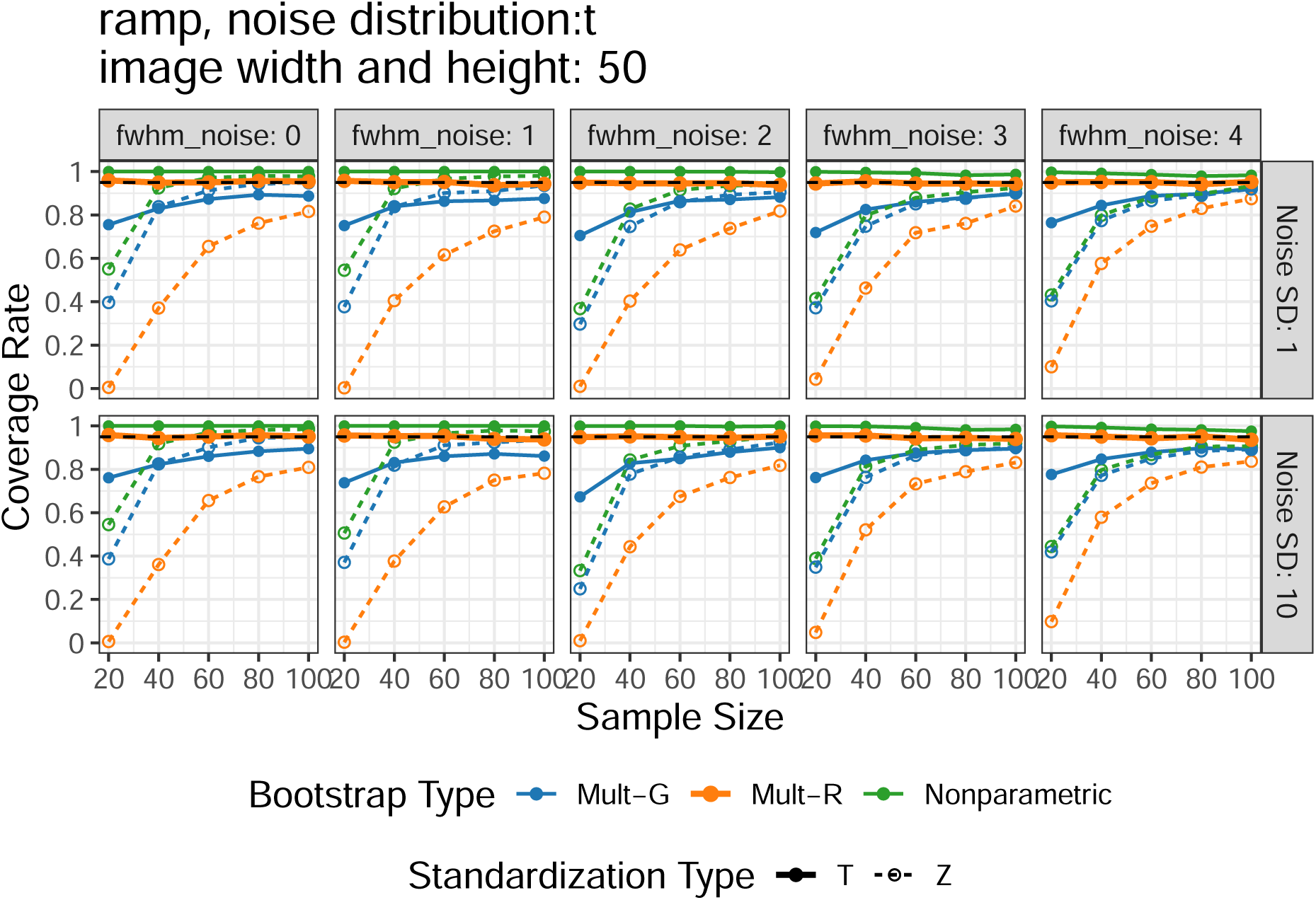

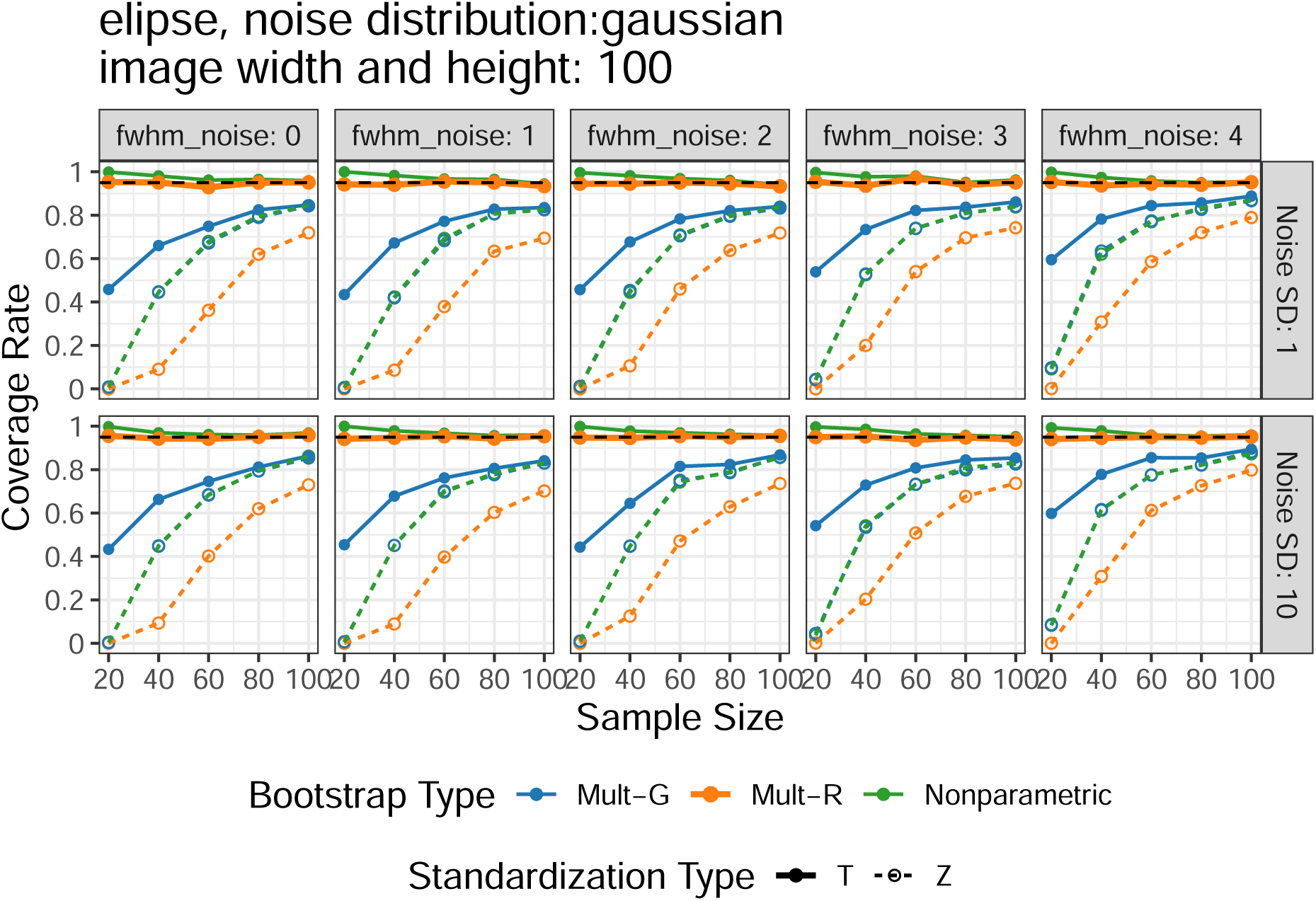

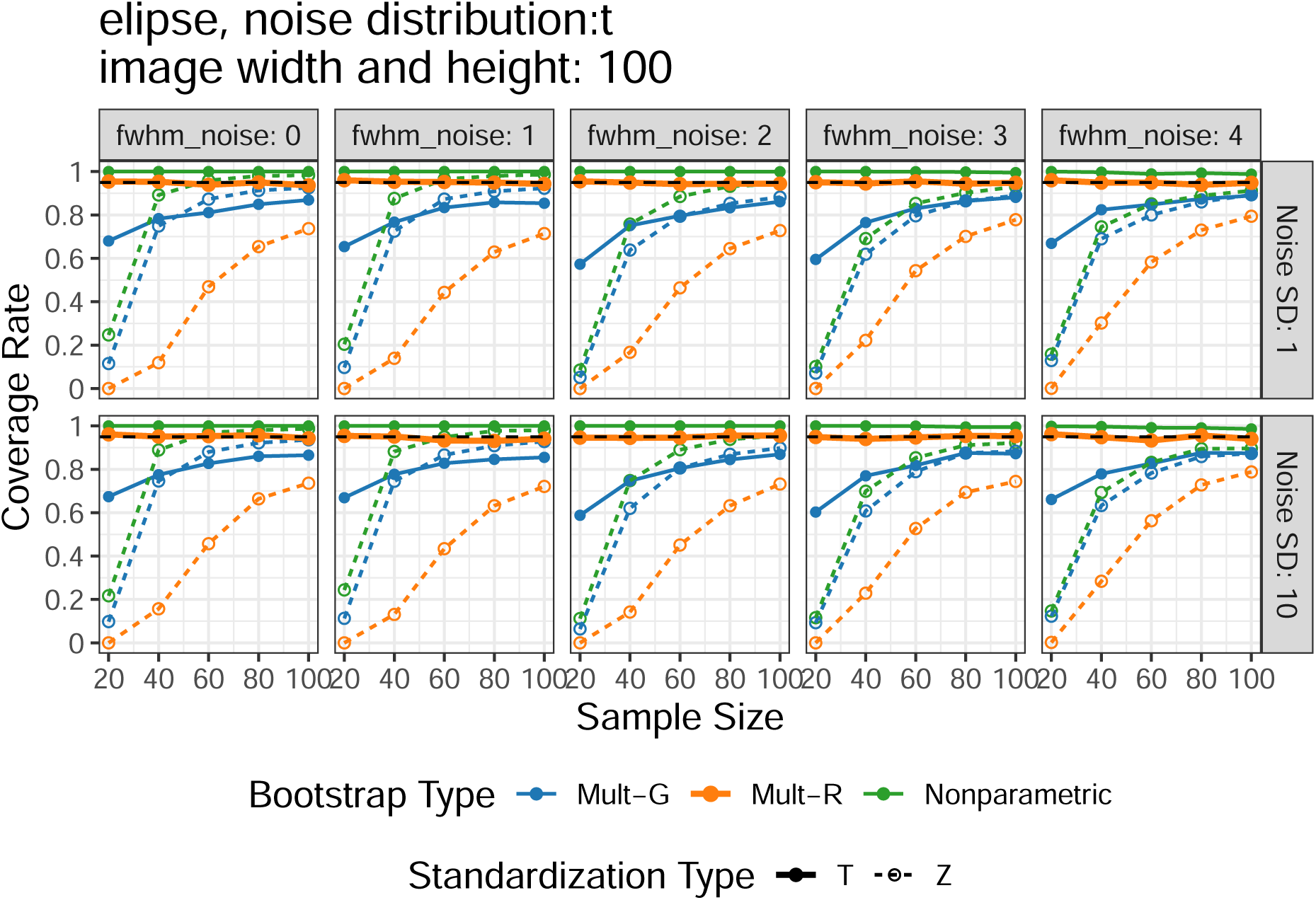

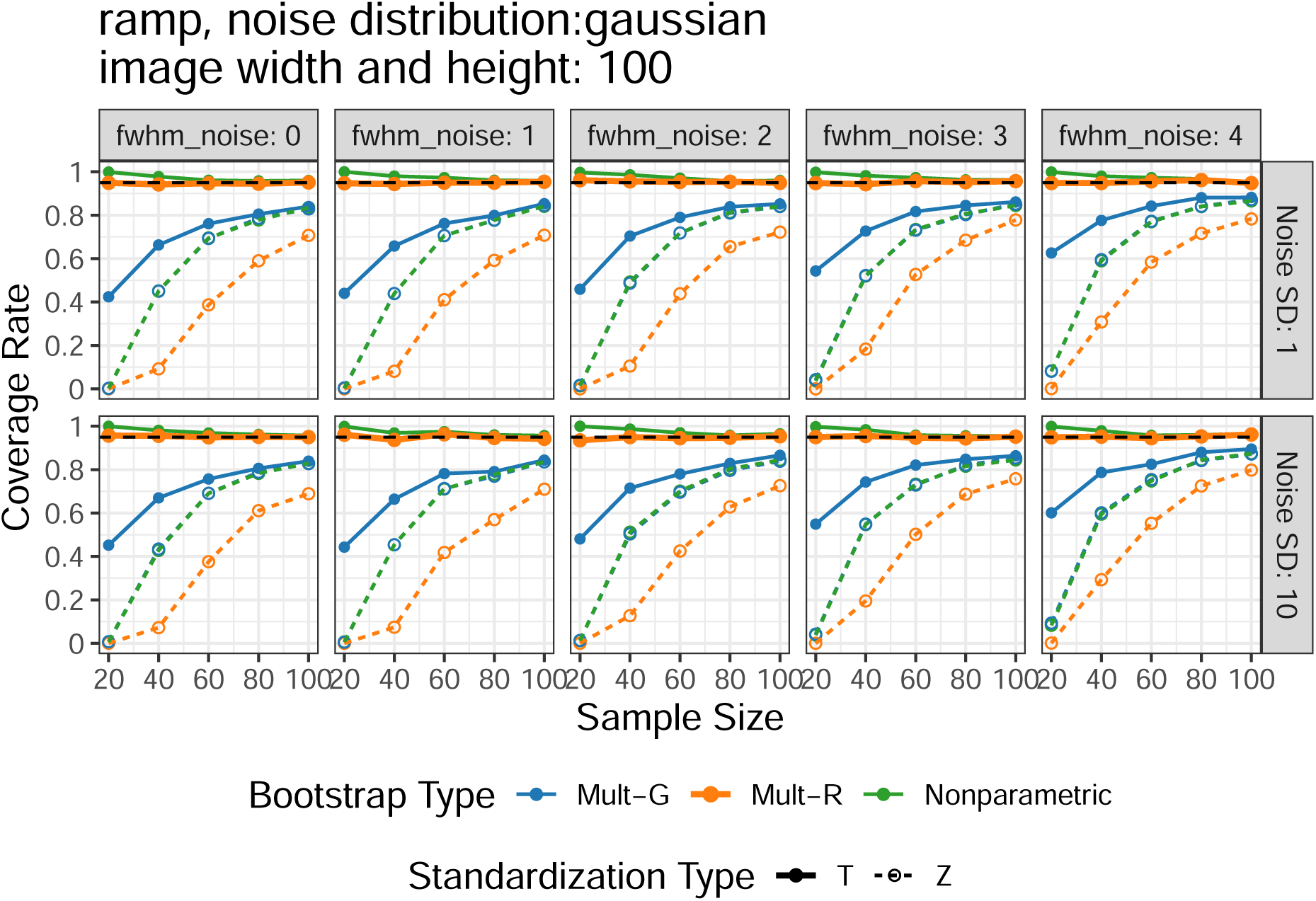

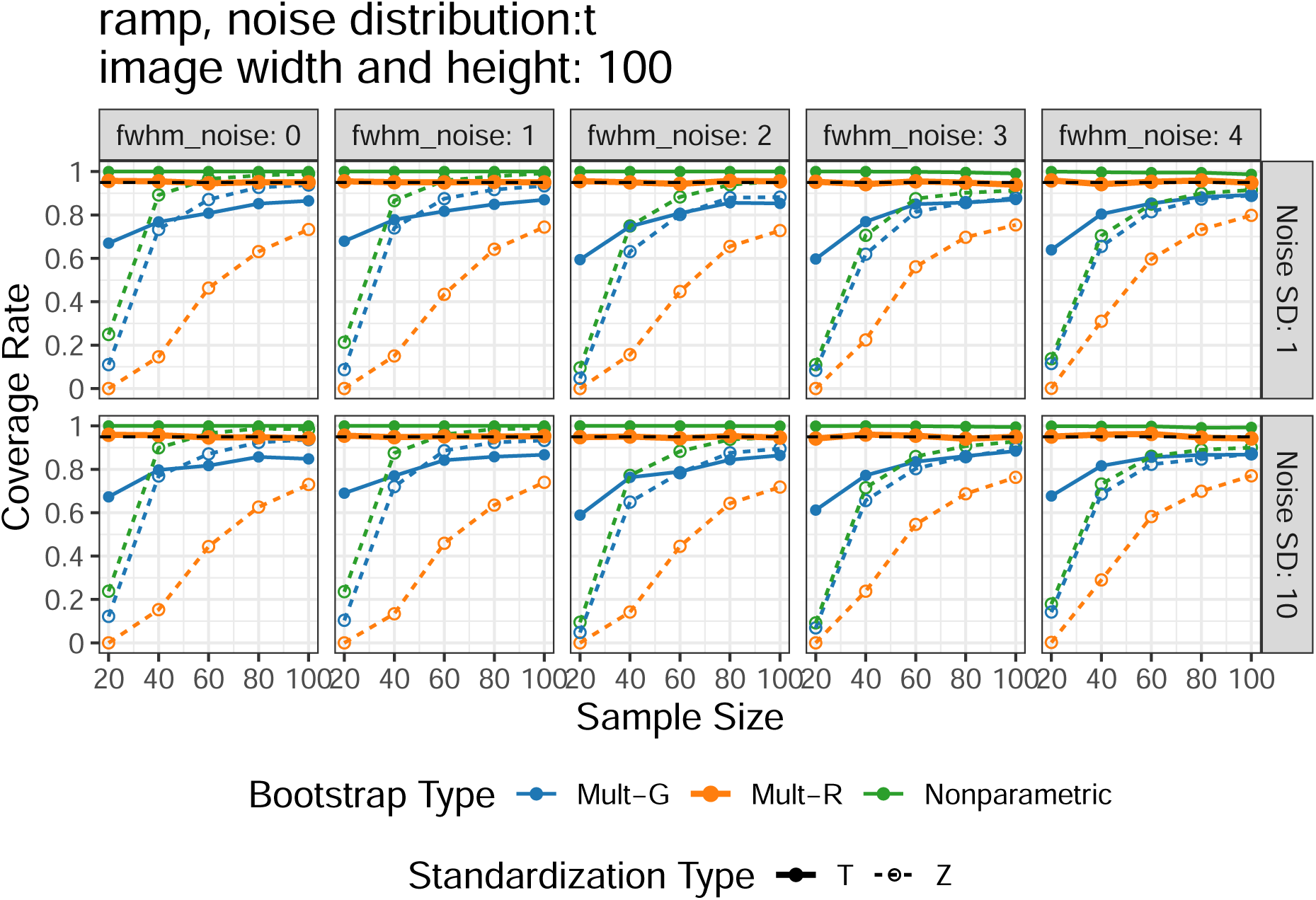

### 2.2 Runtime

Results of runtime in all simulated scenarios:

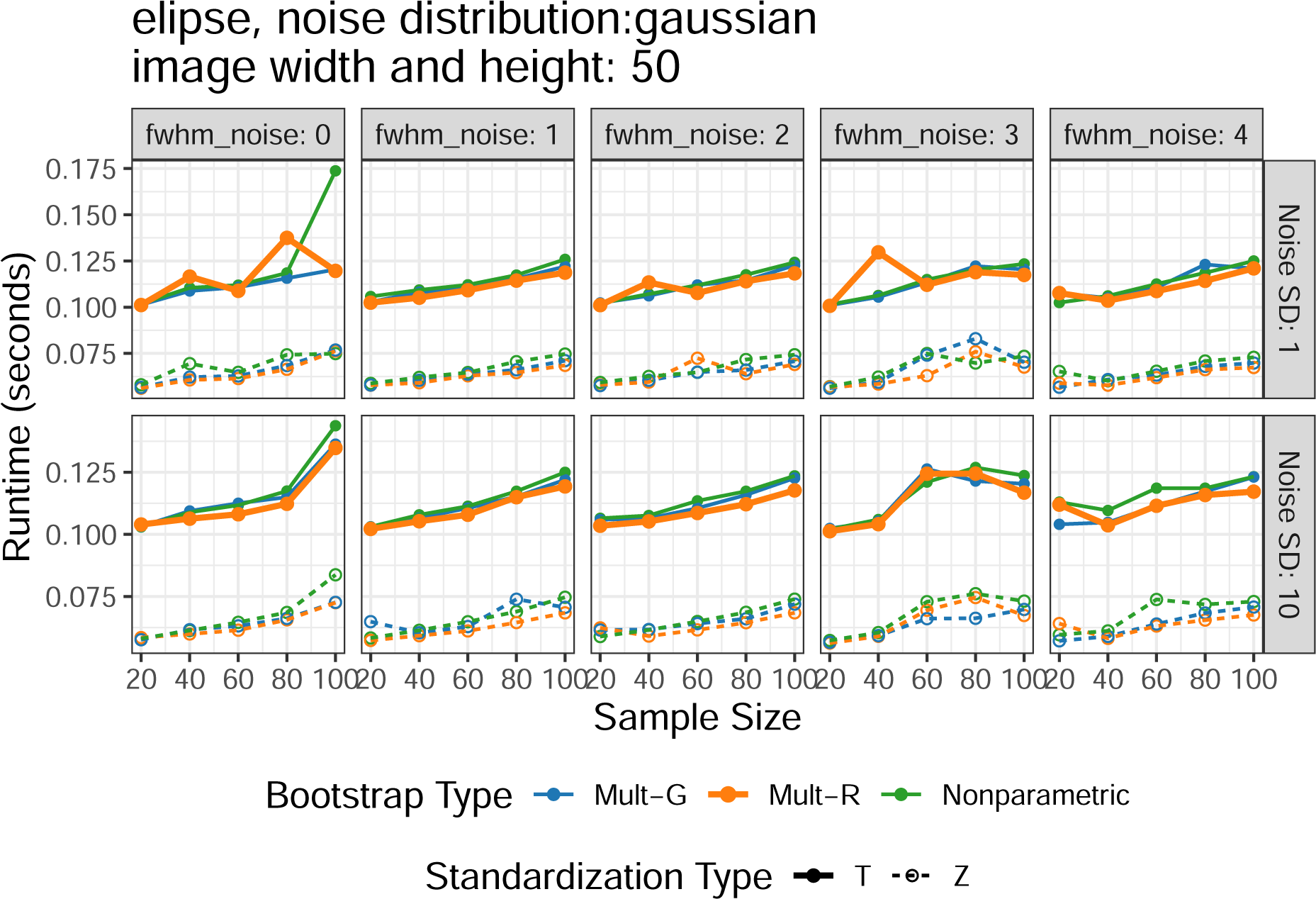

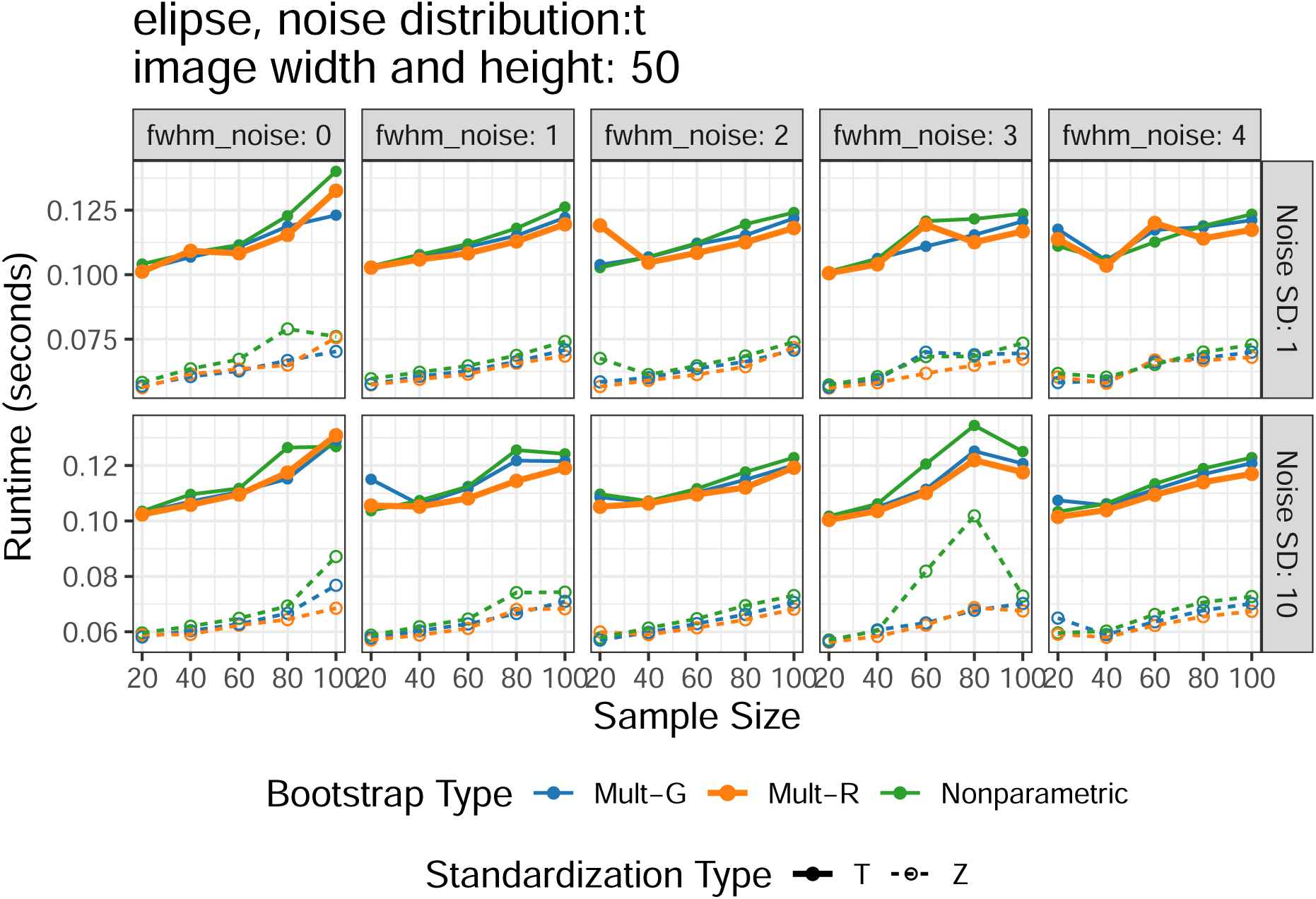

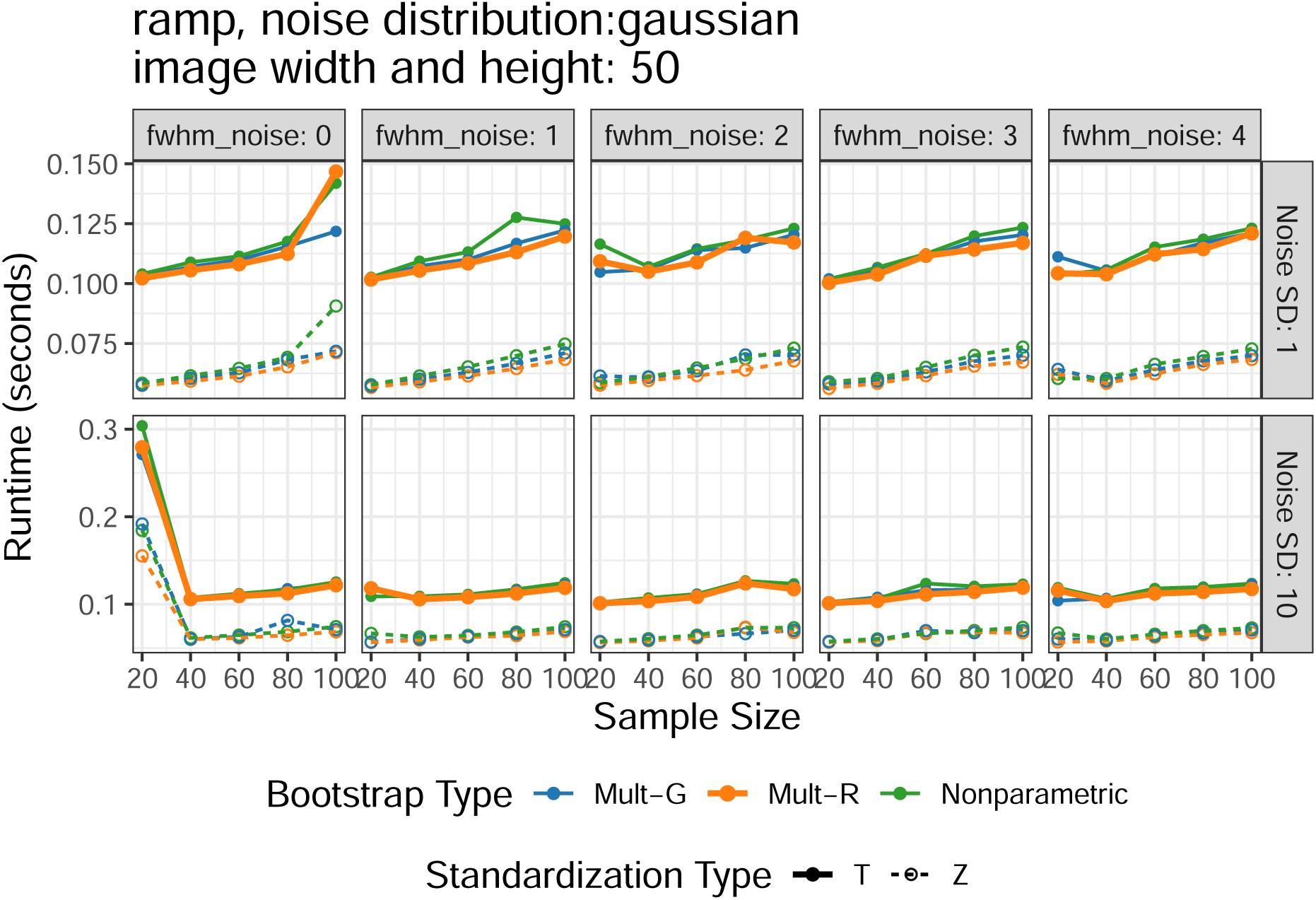

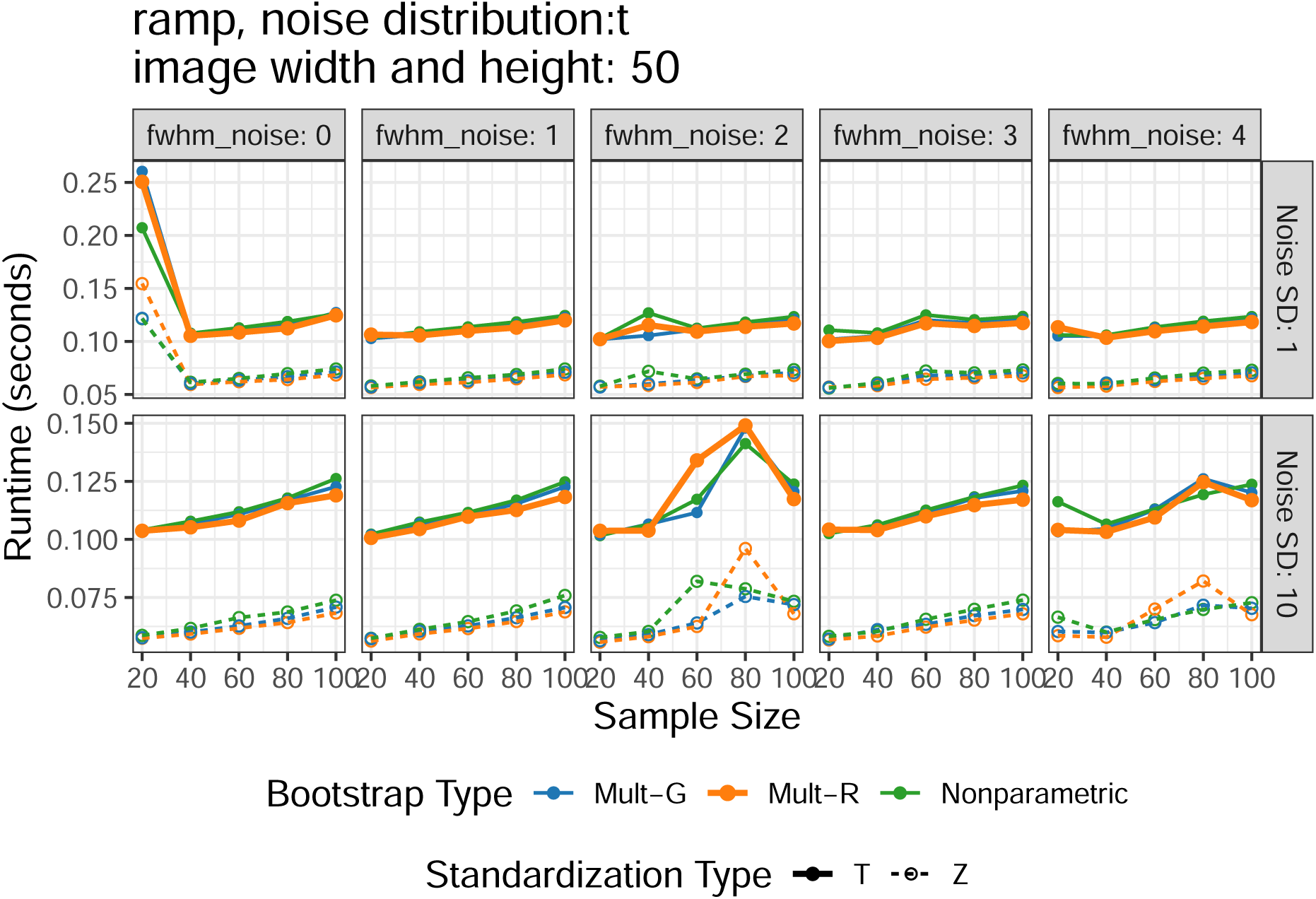

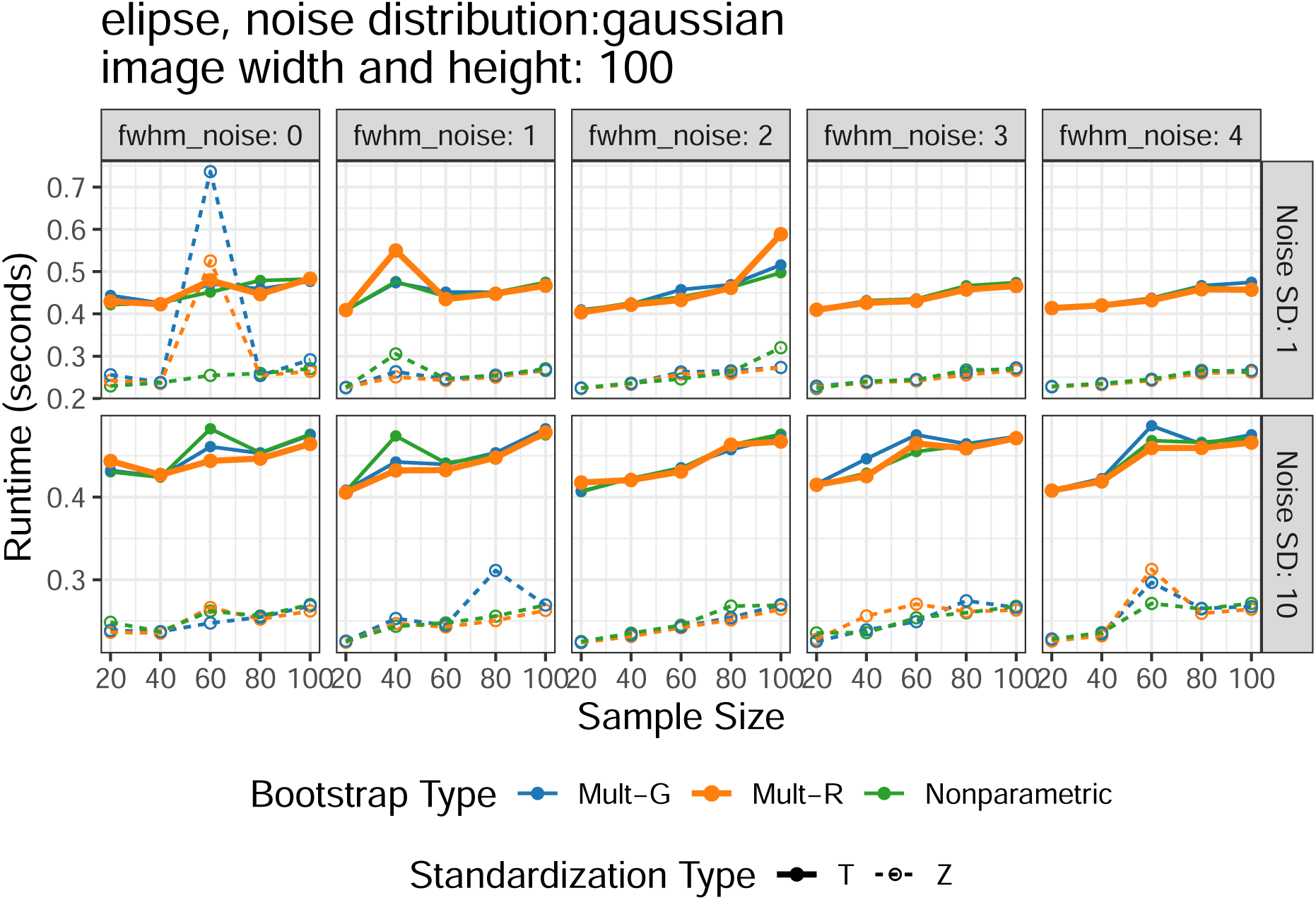

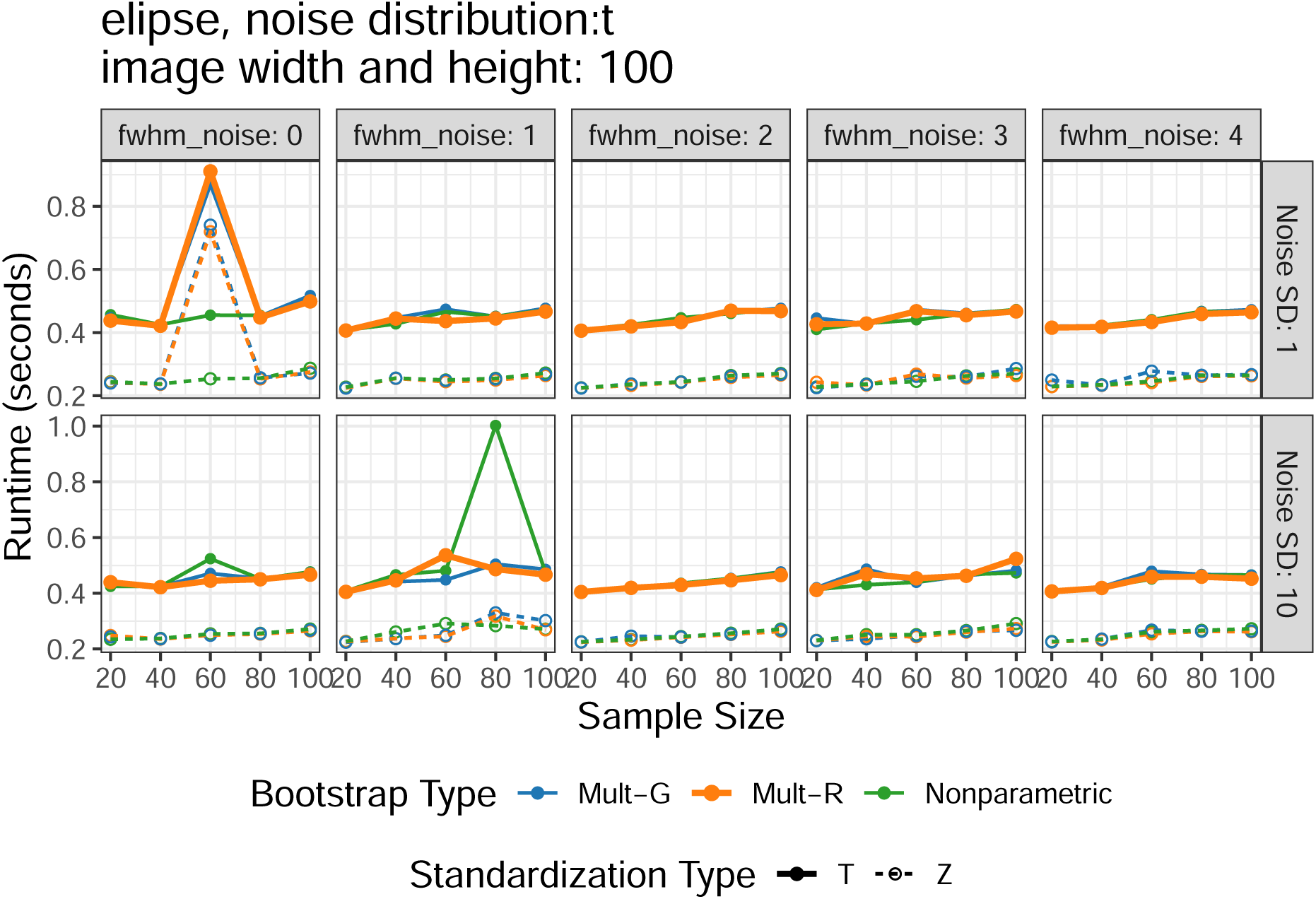

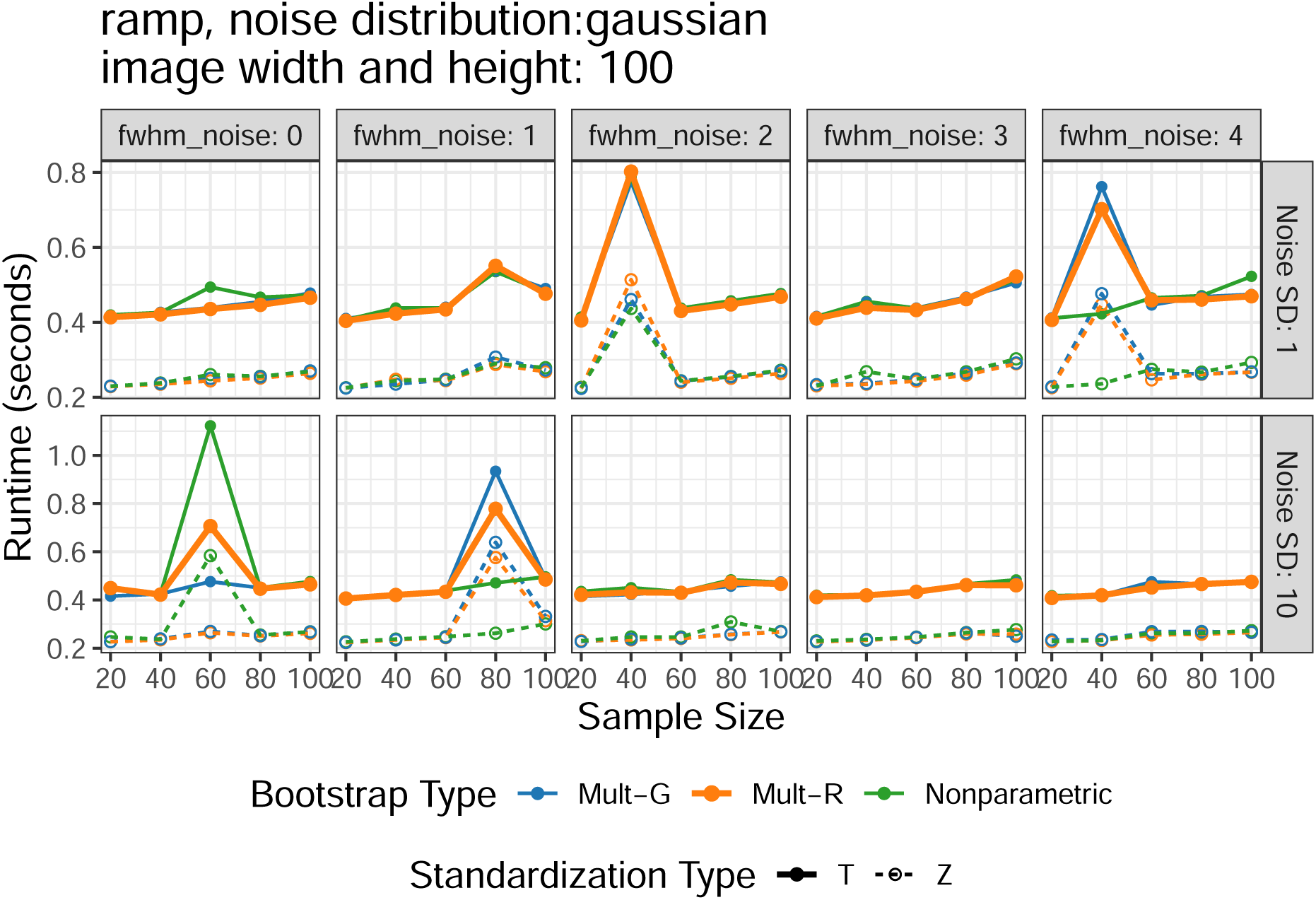

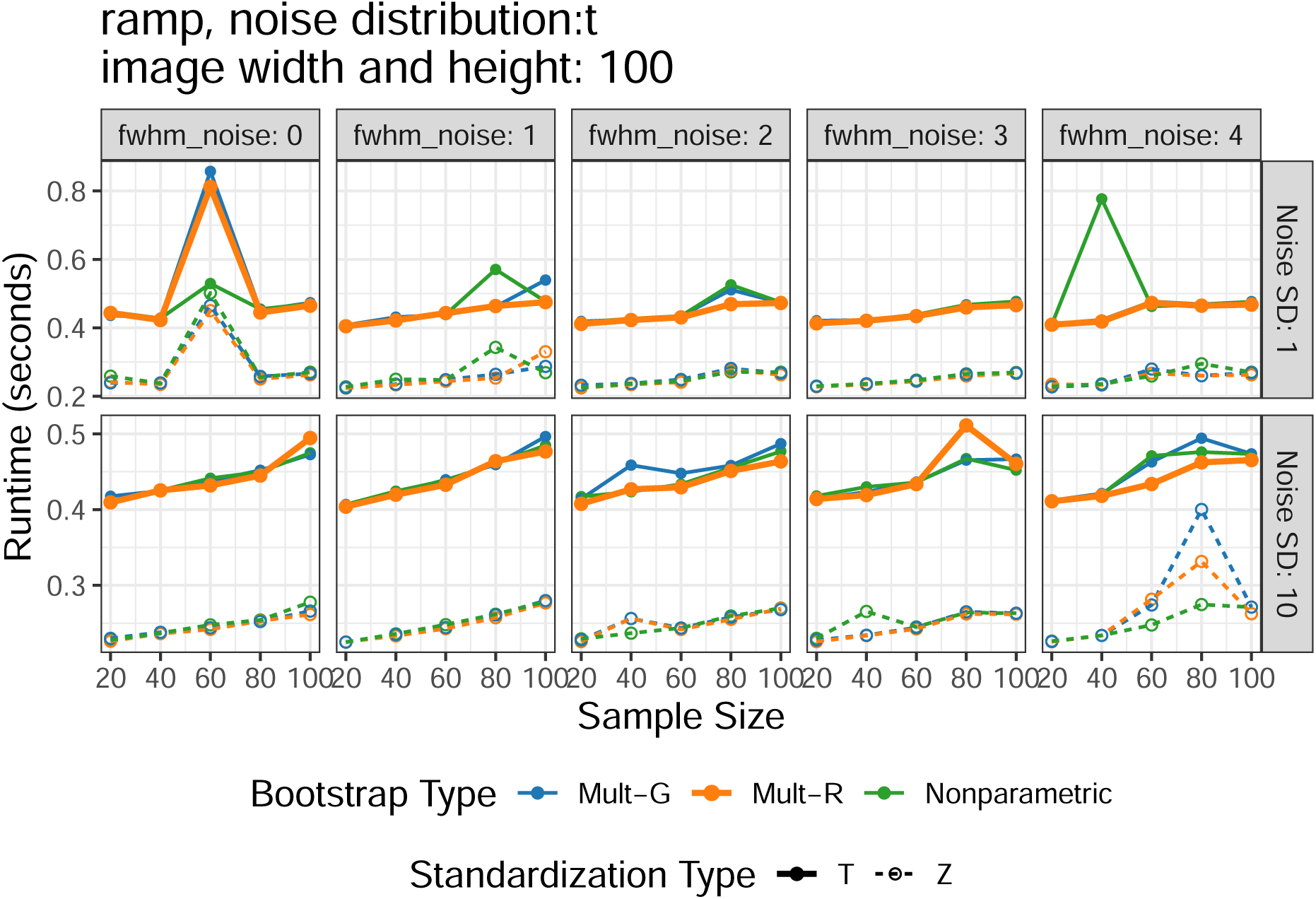

### 2.3 Precision

The plots below are precision results in all simulated scenarios, where a smaller mean quantile represents a more precise SCB. Of note, only methods achieving a relatively good coverage rate were shown here since precision is irrelevant for methods with poor coverage.

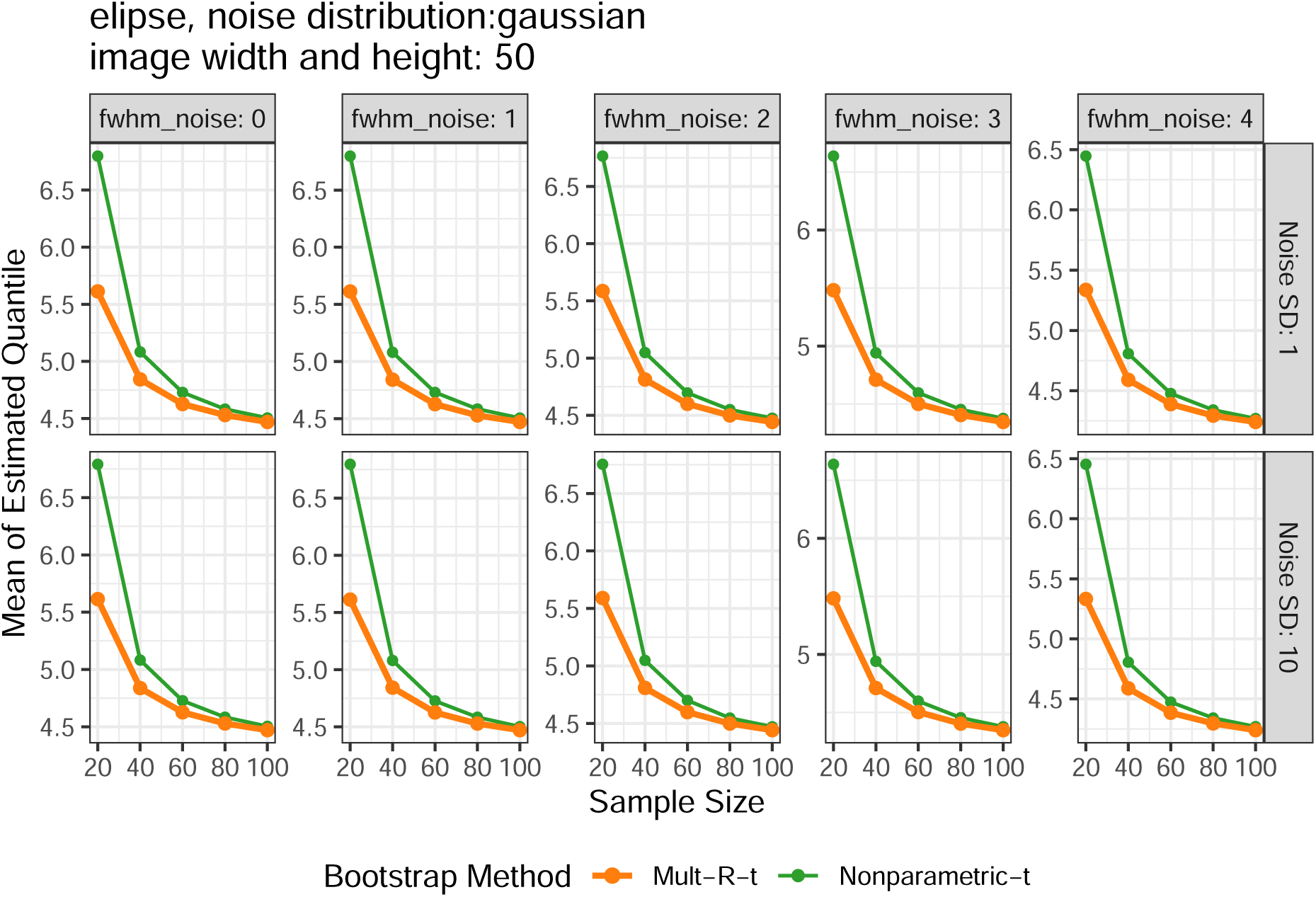

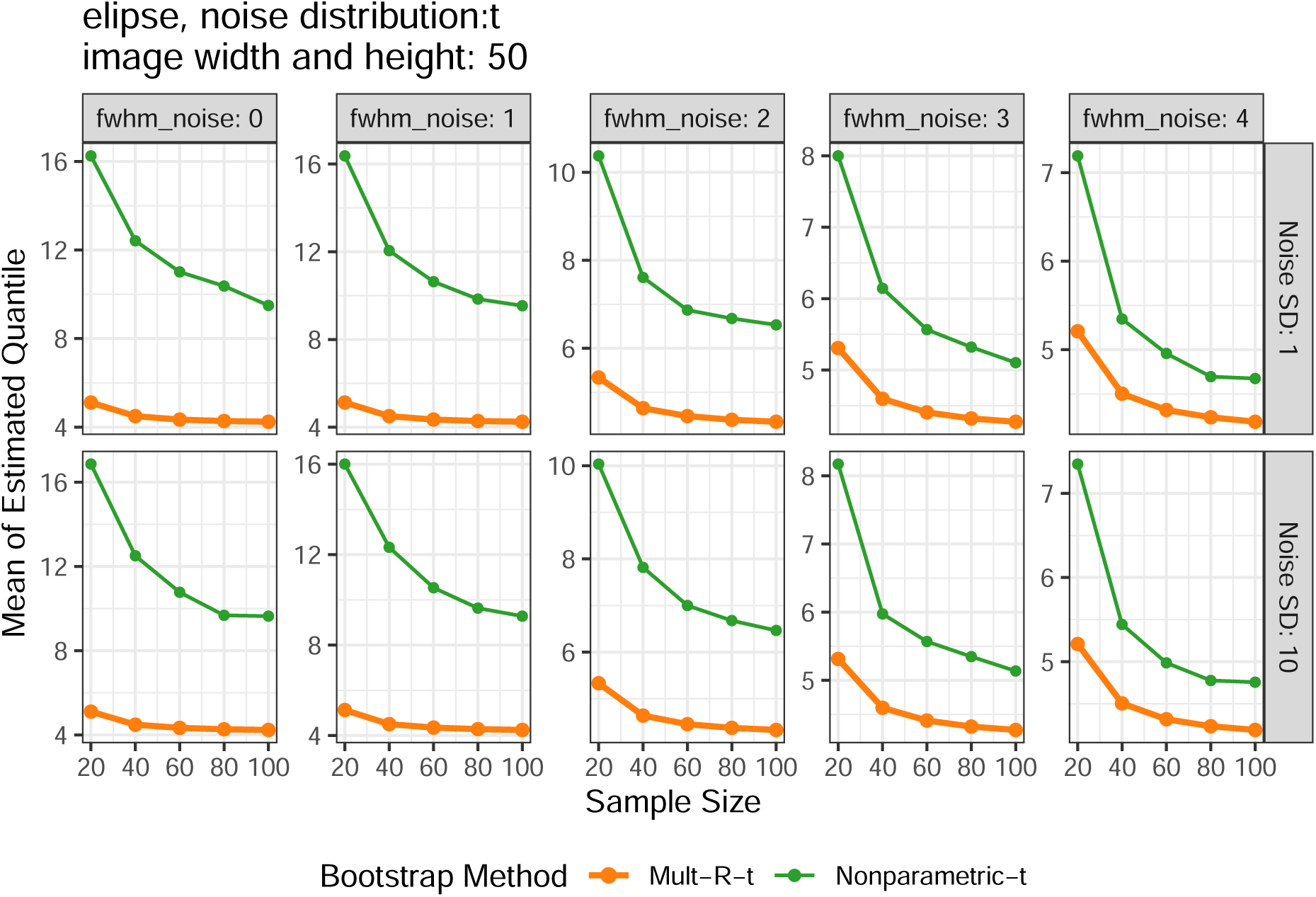

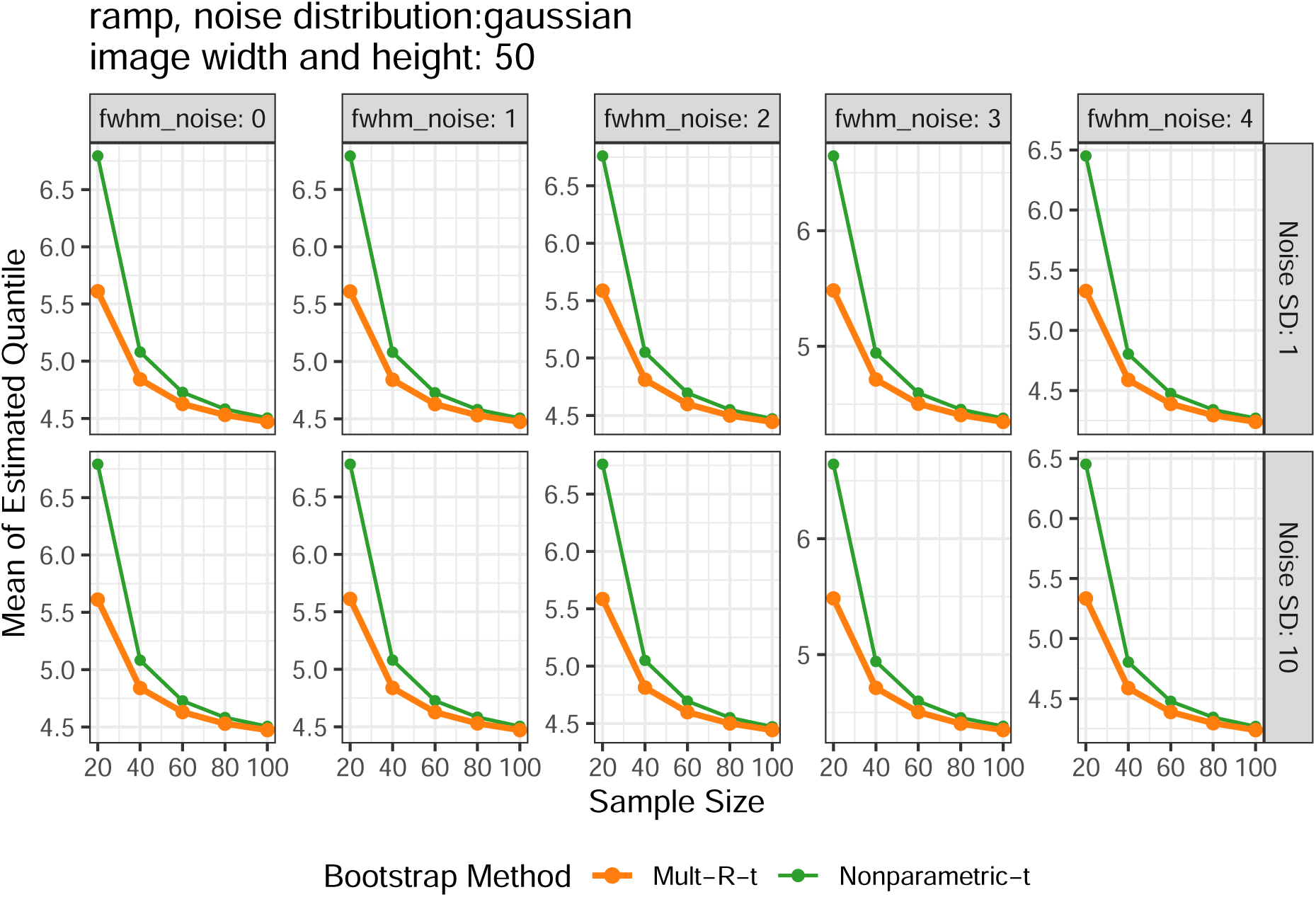

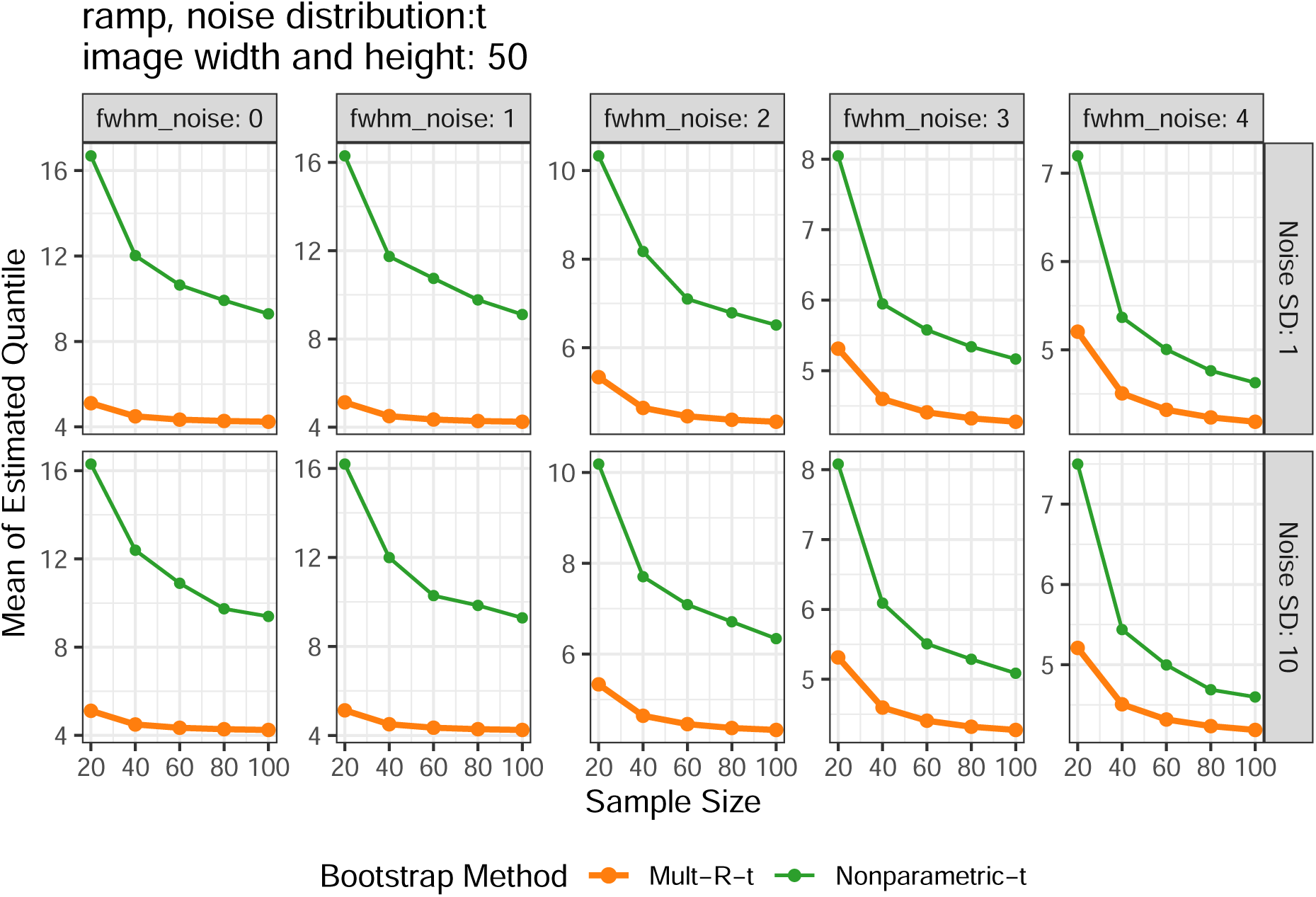

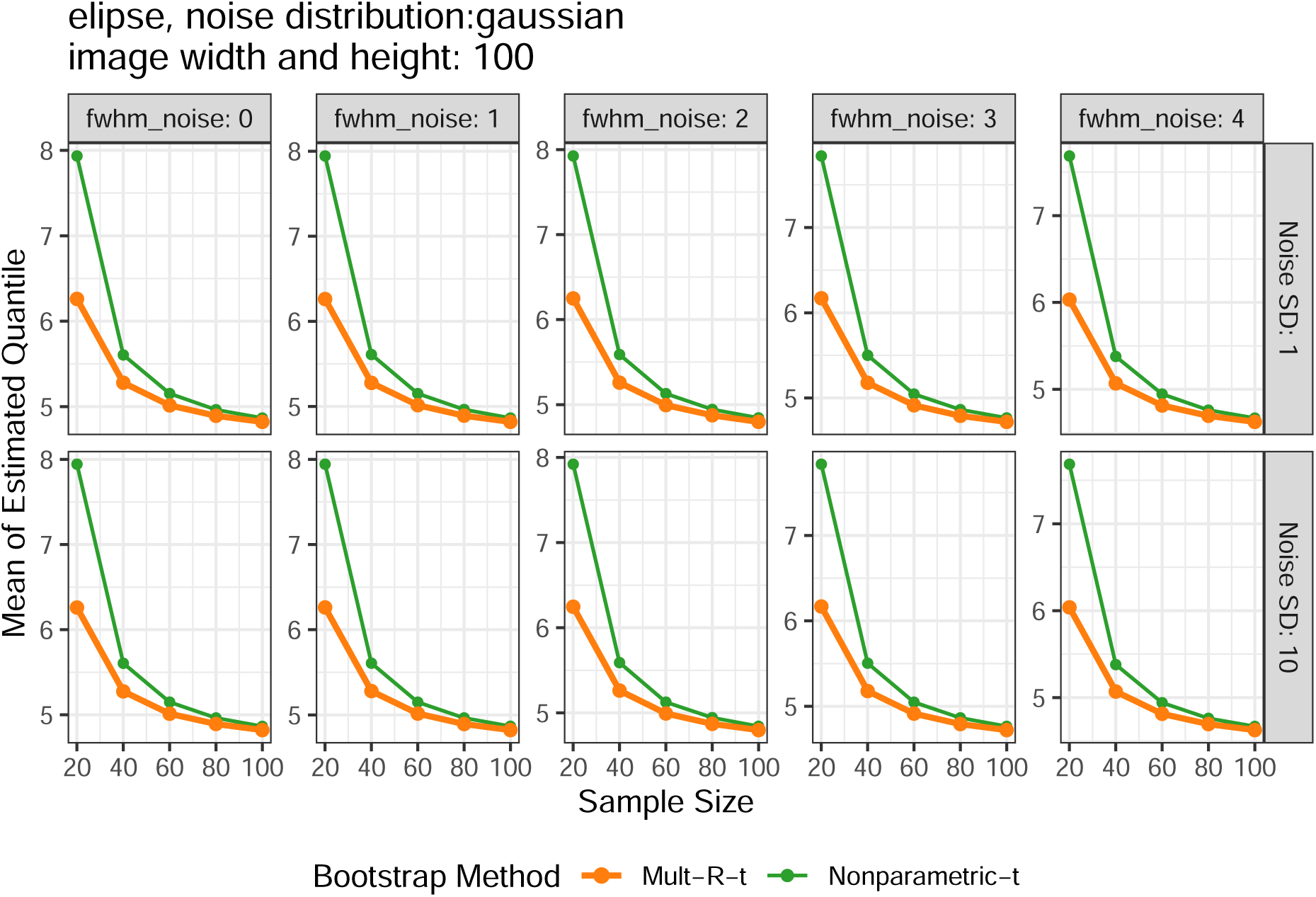

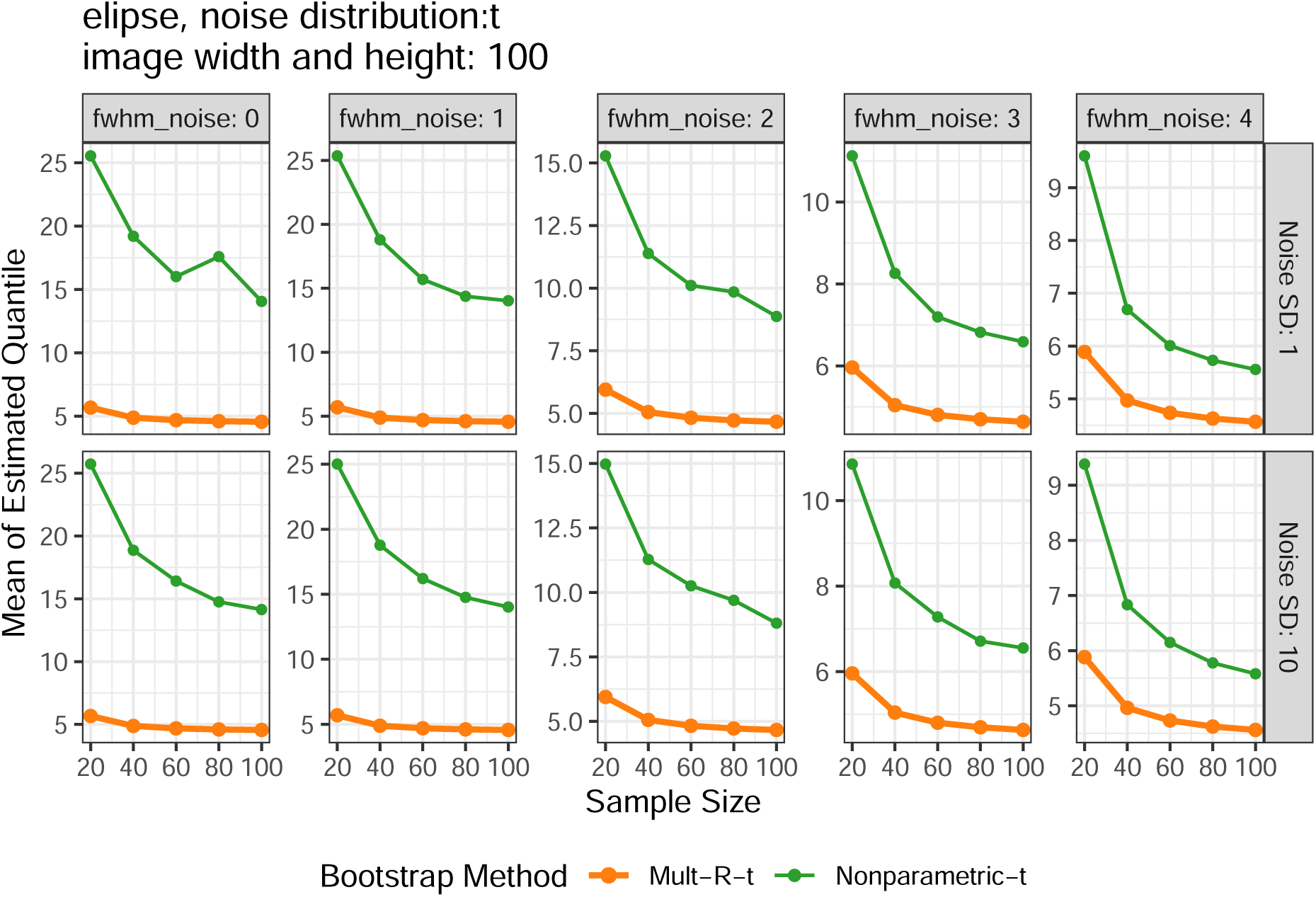

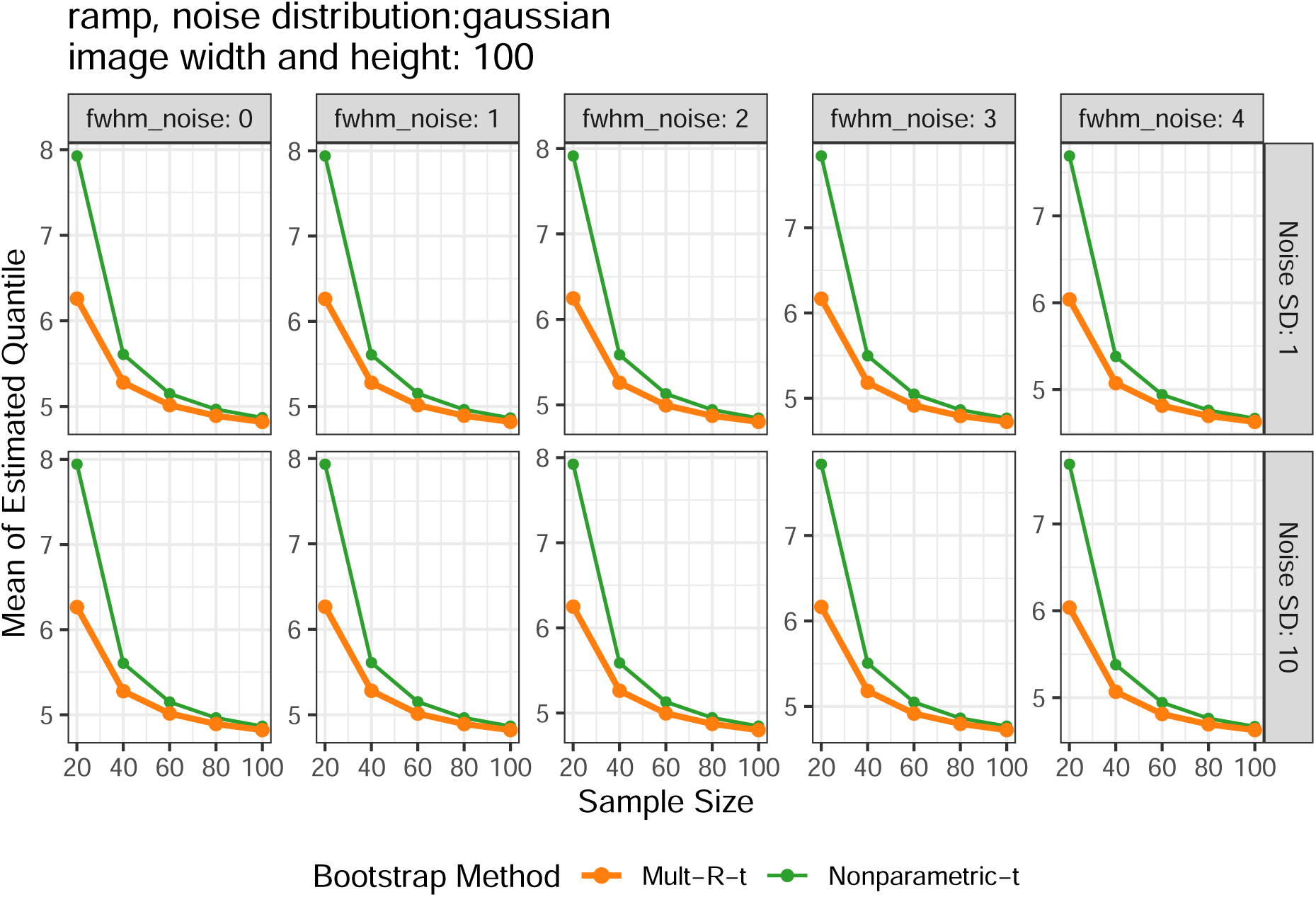

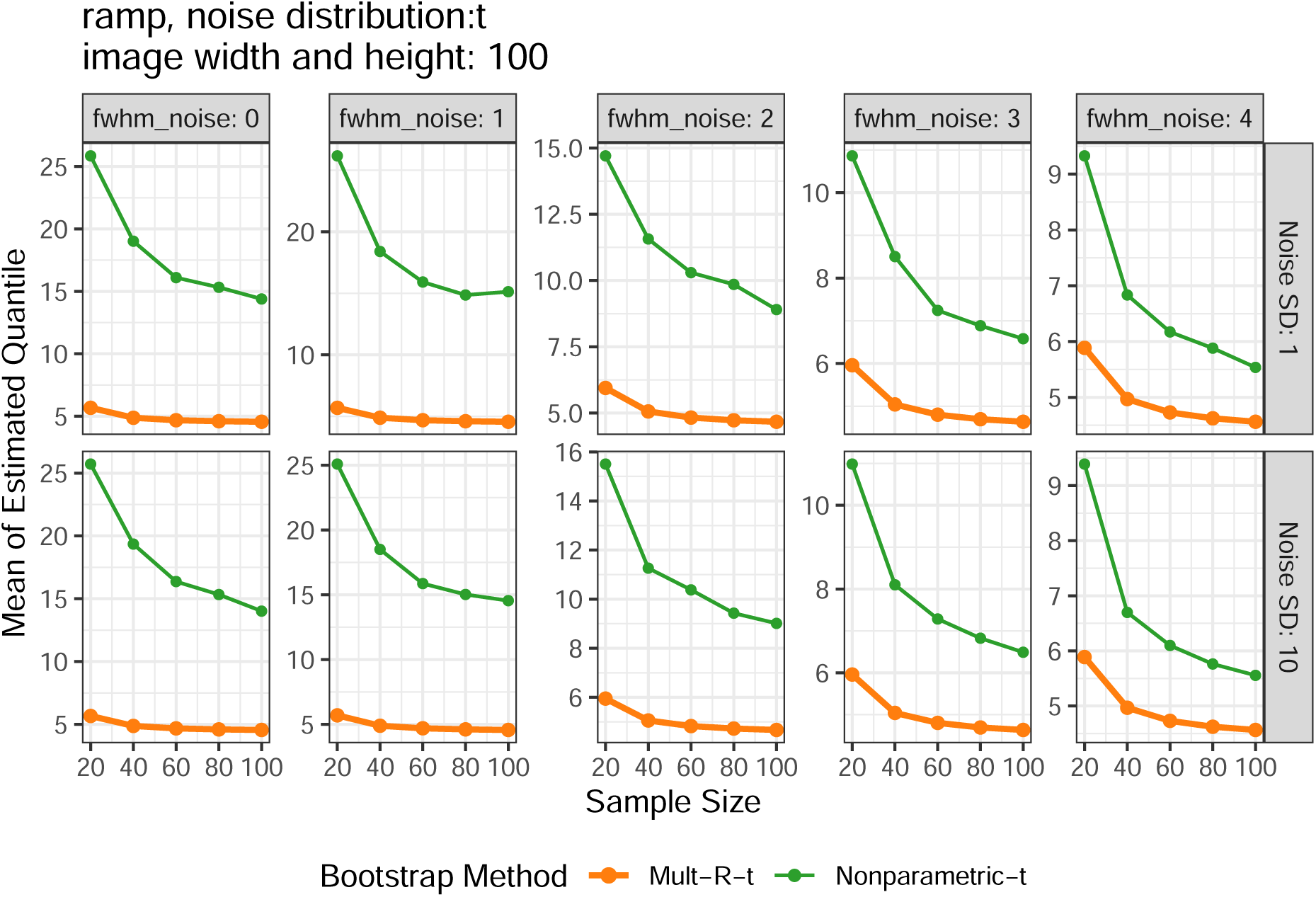

### 2.4 Stability

The plots below are stability results in all simulated scenarios, where a smaller SD of quantile represents a more stable SCB. Of note, only methods achieving a relatively good coverage rate were shown here since stability is irrelevant for methods with poor coverage.

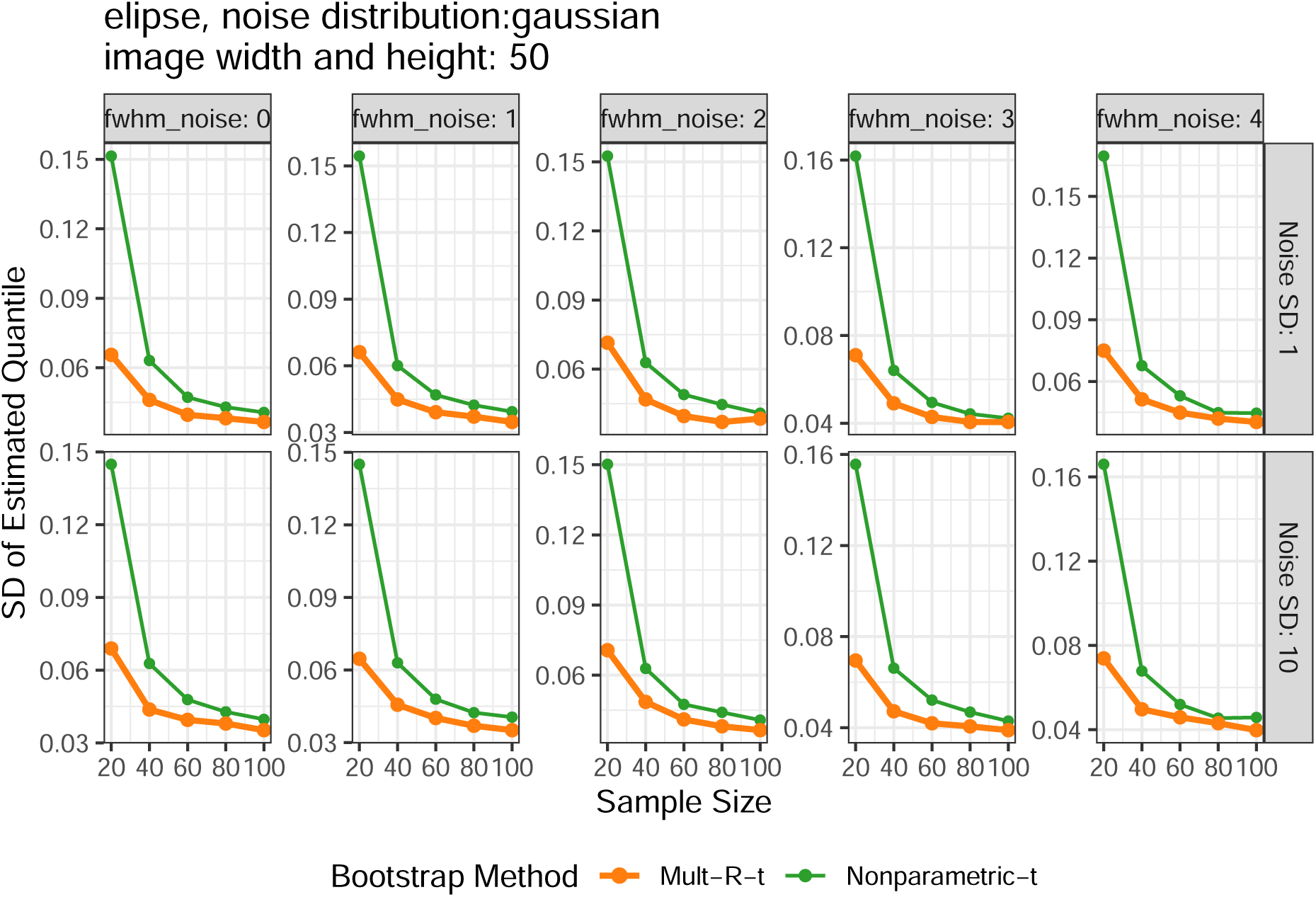

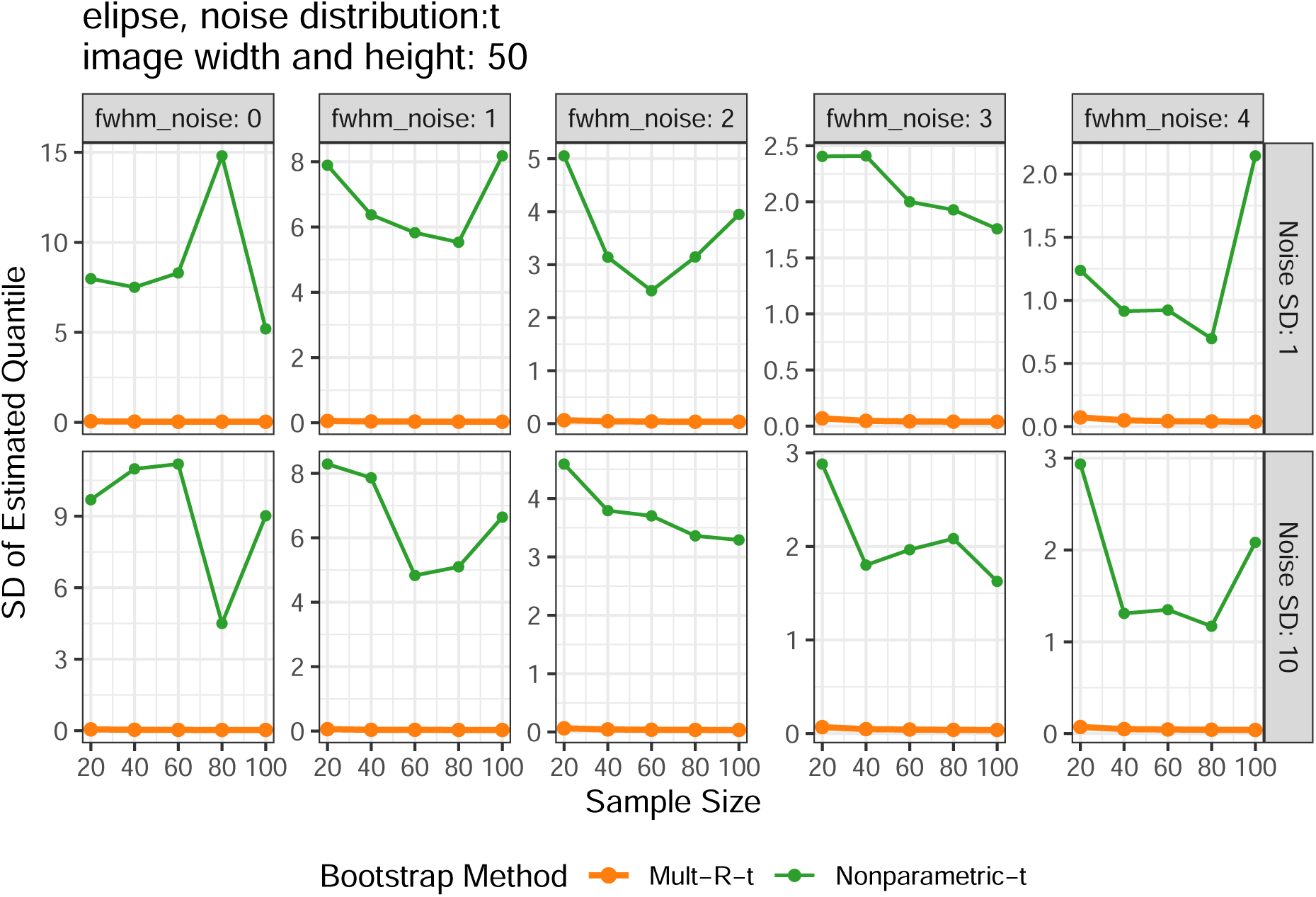

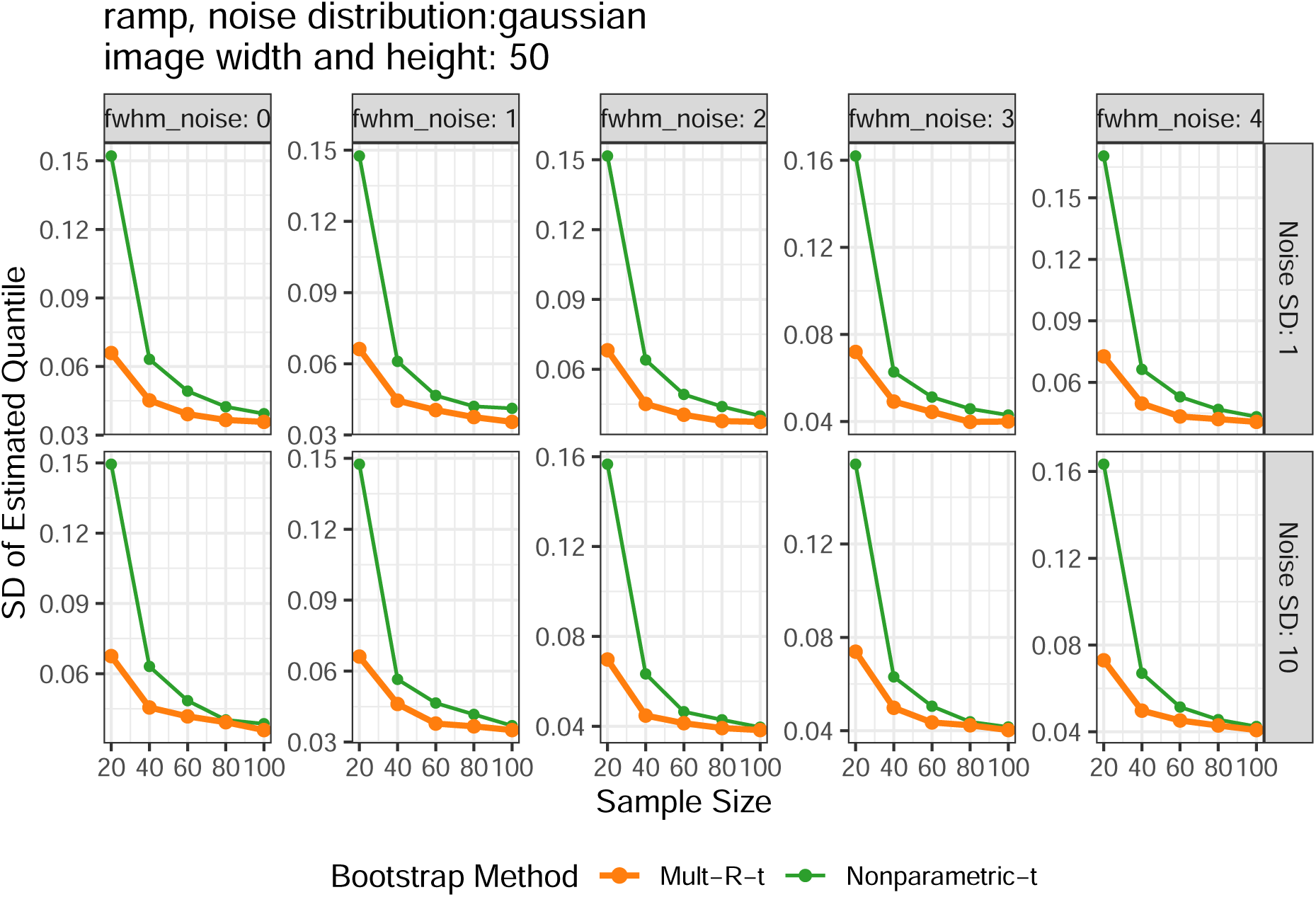

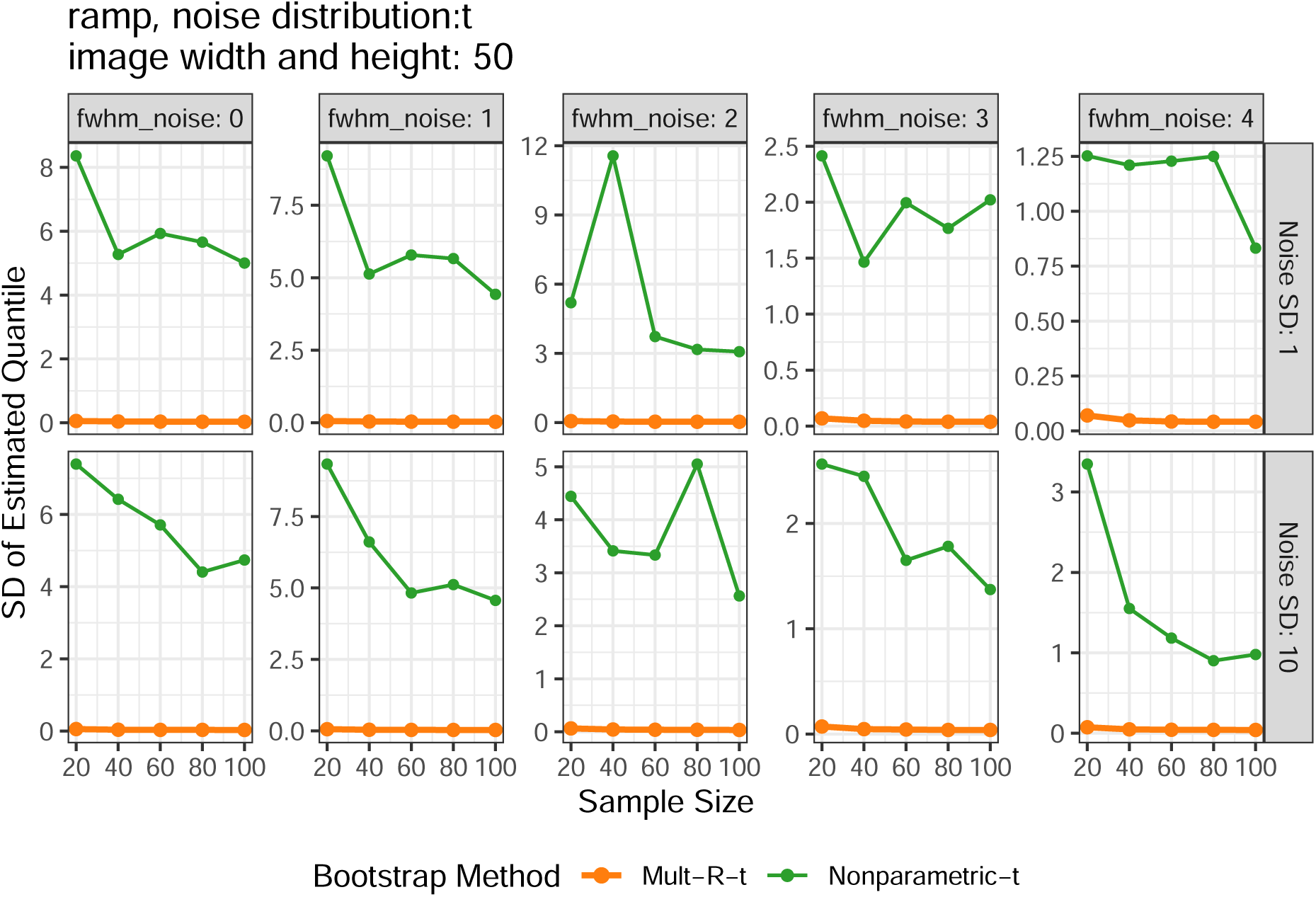

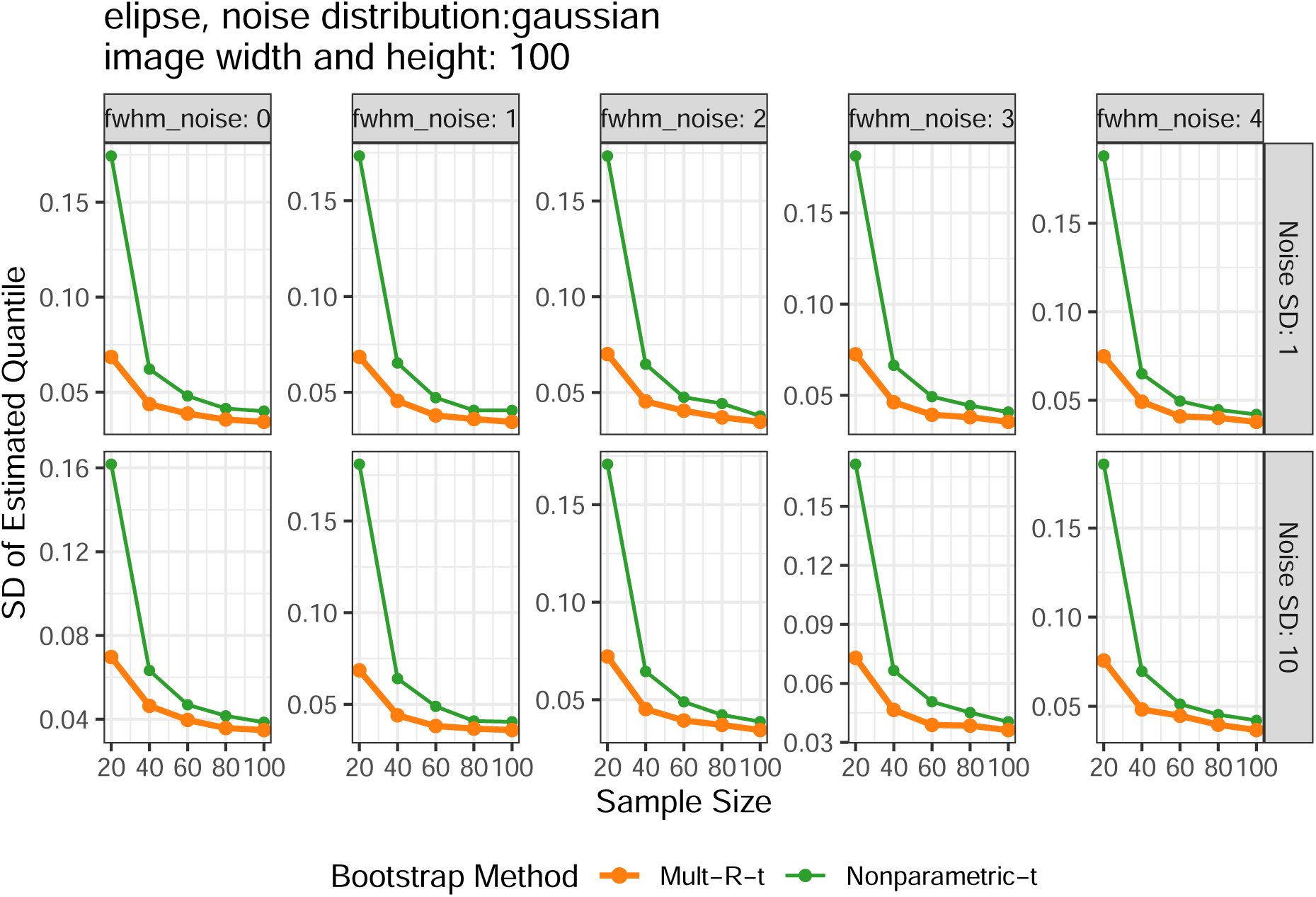

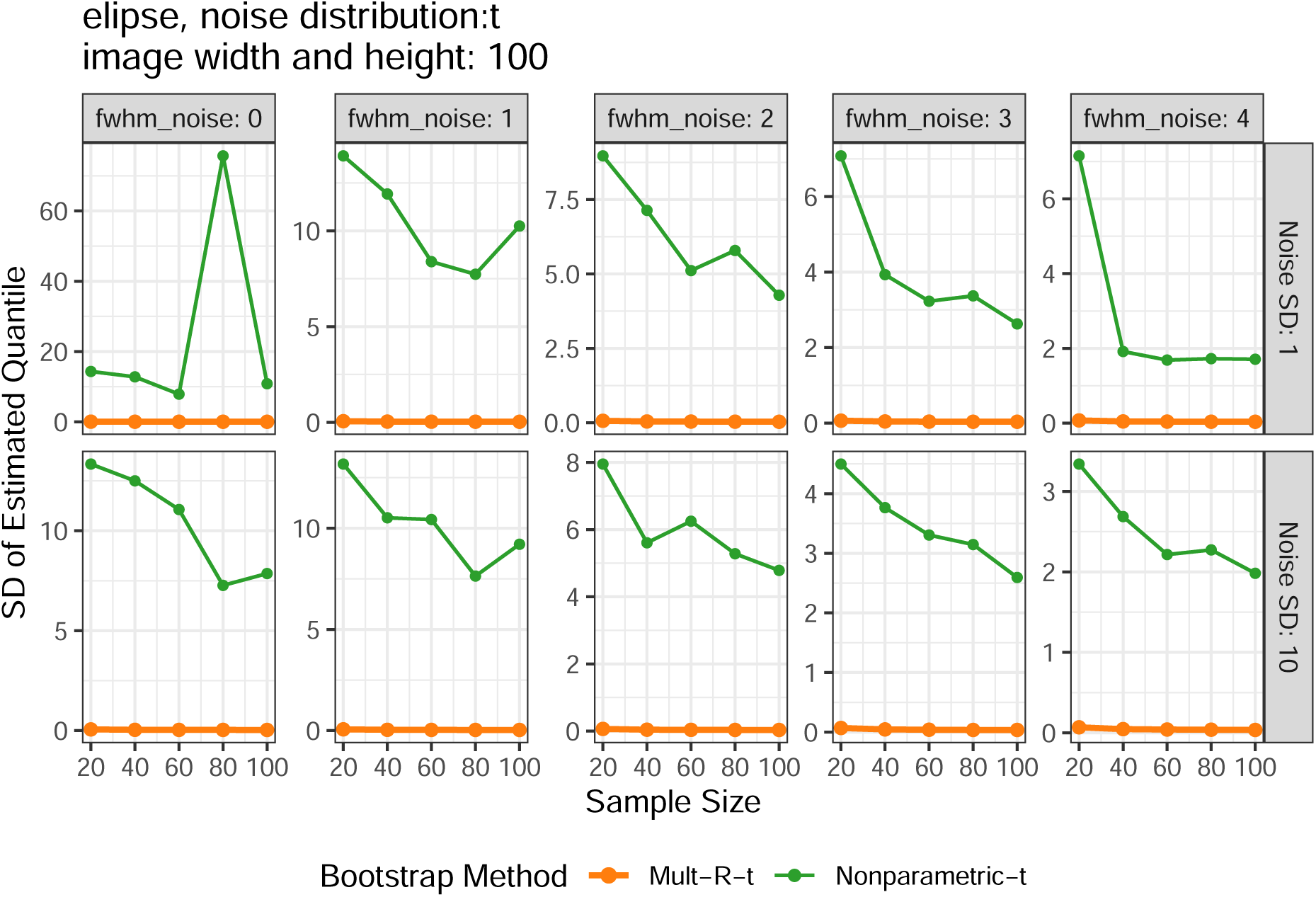

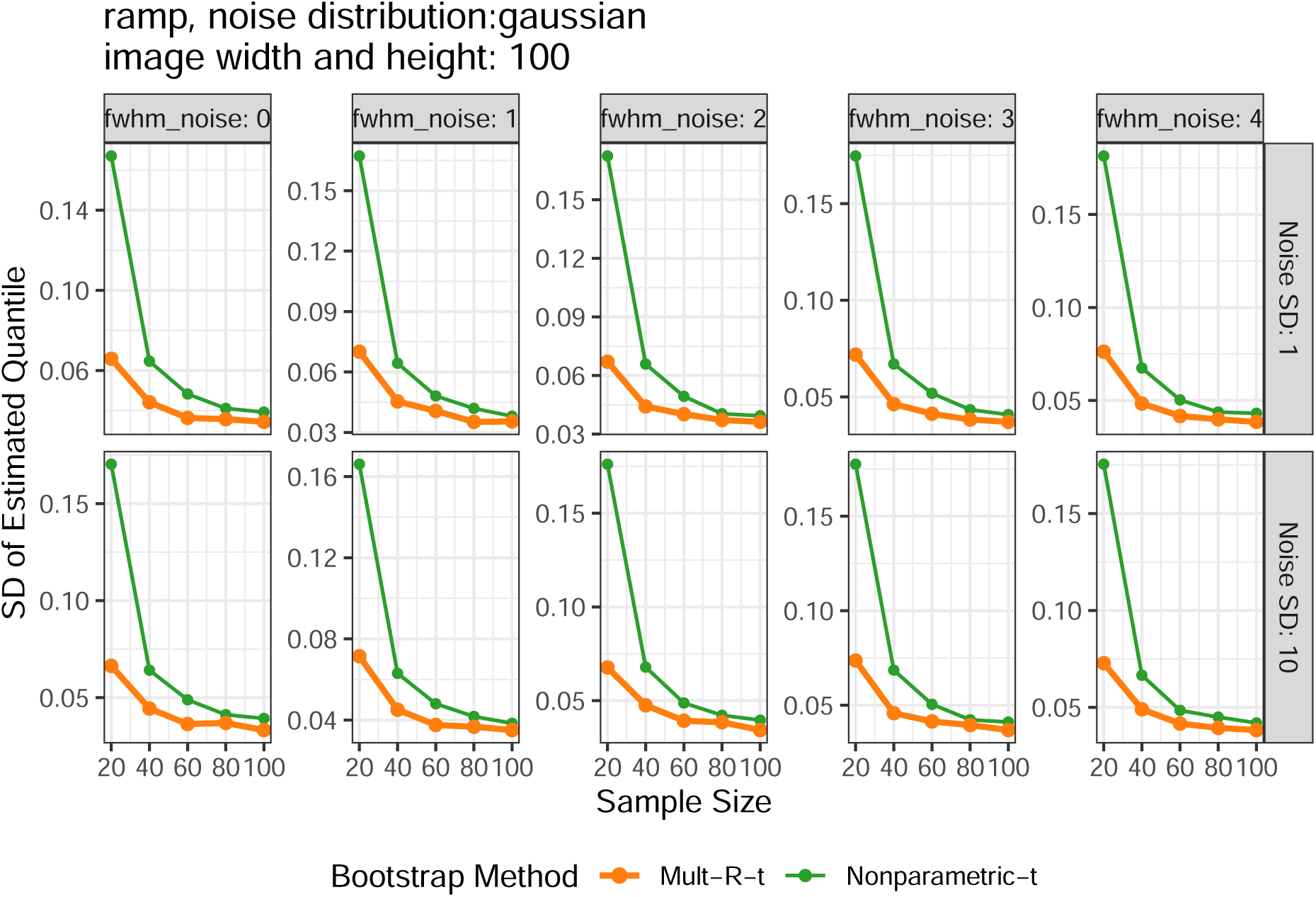

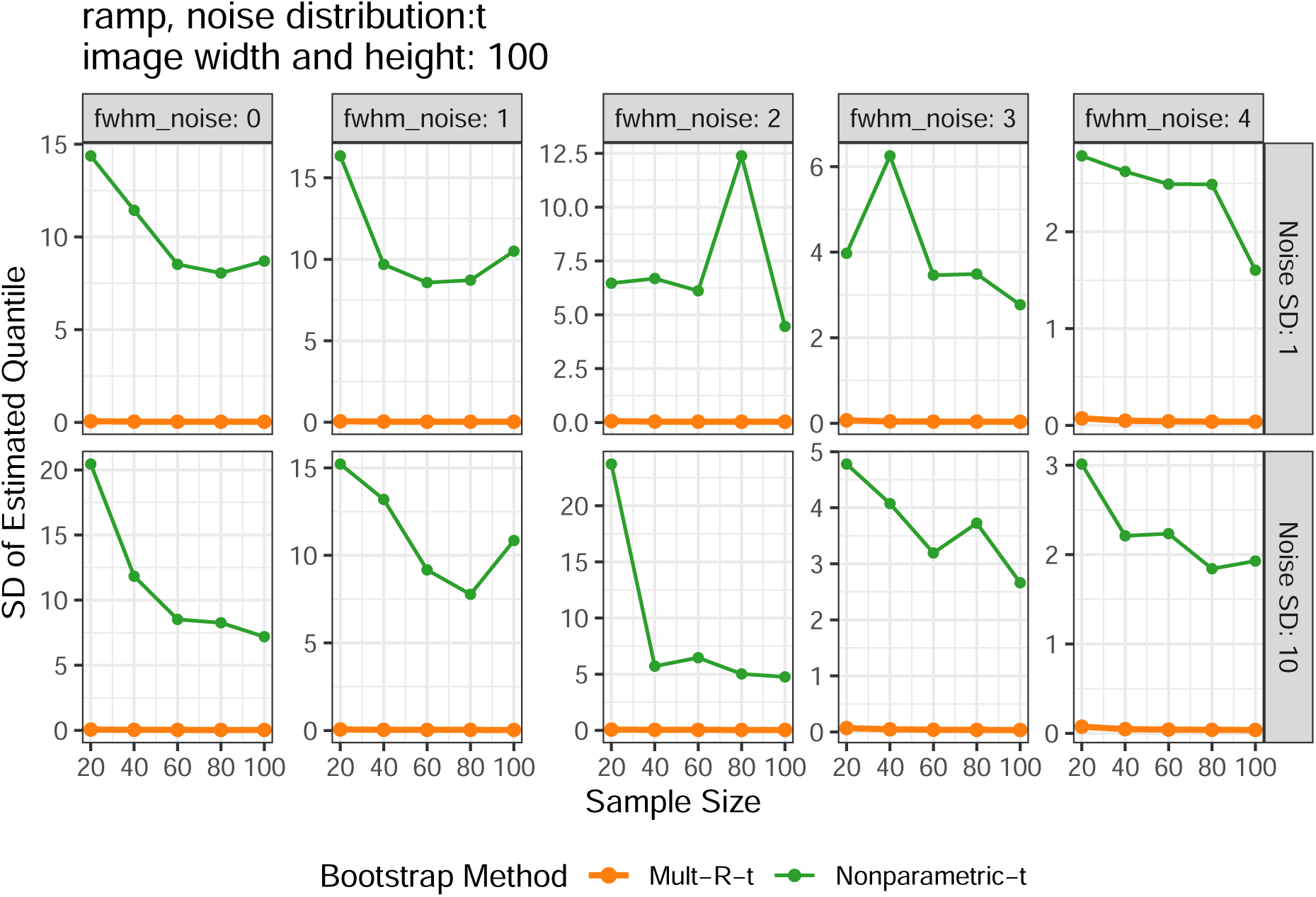

